# Anterior insula-associated social novelty recognition: orchestrated regulation by a local retinoic acid cascade and oxytocin signaling

**DOI:** 10.1101/2021.01.15.426848

**Authors:** Sun-Hong Kim, Kyongman An, Ho Namkung, Matthew D. Rannals, James R. Moore, Tyler Cash-Padgett, Marina Mihaljevic, Sneha Saha, Lina S. Oh, Mari A. Kondo, Kun Yang, Brady J. Maher, Minae Niwa, Akira Sawa

## Abstract

**Background:** Deficits in social cognition consistently underlie functional disabilities in a wide range of psychiatric disorders. Neuroimaging studies have suggested that the anterior insula is a ‘common core’ brain region that is impaired across neurological and psychiatric disorders, which include social cognition deficits. Nevertheless, neurobiological mechanisms of the anterior insula for social cognition remain elusive.

**Methods:** To determine the role of anterior insula in social cognition, we manipulated expression of Cyp26B1, an anterior insula-enriched molecule that is crucial for retinoic acid degradation and involved in the pathology of neuropsychiatric conditions. Social cognition was mainly assayed using the three-chamber social interaction test. We conducted multimodal analyses at the molecular, cellular, circuitry, and behavioral levels.

**Results:** At the molecular/cellular level, anterior insula-mediated social novelty recognition is maintained by proper activity of the layer 5 pyramidal neurons, for which retinoic acid-mediated gene transcription can play a role. We also demonstrate that oxytocin influences the anterior insula-mediated social novelty recognition, not by direct projection of oxytocin neurons, nor by direct diffusion of oxytocin to the anterior insula, which contrasts the modes of oxytocin regulation onto the posterior insula. Instead, oxytocin affects oxytocin receptor-expressing neurons in the dorsal raphe nucleus where serotonergic neurons are projected to the anterior insula. Furthermore, we show that serotonin 5HT2C receptor expressed in the anterior insula influences social novelty recognition.

**Conclusions:** Anterior insula plays a pivotal role in social novelty recognition that is partly regulated by a local retinoic acid cascade, but also remotely regulated by oxytocin via a non-classic mechanism.

## Introduction

Social cognition is critical for human behavior in complex social environments (1, 2). Accordingly, deficits in social cognition consistently underlie functional disabilities in individuals suffering from a wide range of psychiatric disorders (3–7). Social novelty recognition (the ability to recognize social cues), a key subdomain of social cognition, can be assessed by performance-based measures of social cue (e.g. face or voice perception) in humans (8, 9) as well as by social novelty recognition/preference paradigms (3-chamber social interaction test and 5-trial social novelty recognition test) in rodents (10, 11).

The insular cortex was one of the least understood brain regions until recently (12). However, the advent of functional imaging techniques have recently unveiled its multiple functions in humans, which include interoception, emotion, cognition, and motivation (13–15). These studies, including a recent meta-analysis, have suggested that the insular cortex is a ‘common core’ brain region that is impaired across neurological and psychiatric disorders (2, 13, 16–19). The insular cortex can be divided into the anterior insula (AI) and posterior insula (PI). They are different with each other in the cytoarchitecture, connections, and functions, but they are also tightly interconnected. The PI shows a classical 6-layer structure, whereas the AI lacks granular layer 4 (18). The PI receives multiple sensory inputs (18, 19), whereas the AI makes heavy reciprocal connections with the frontal areas and limbic system and participates in the salience network (19).

Oxytocin (OT) is a well-appreciated neuropeptide that influences social behavior, including social recognition, pair bonding, mating, and parental care (20–22). OT modulates these behaviors by being released into the target brain regions, such as the amygdala, ventral tegmental area, and dorsal raphe, from the axonal fibers of oxytocinergic neurons located in the paraventricular nucleus (PVN) and supraoptic nucleus (SON) (23, 24). Animal studies for key molecules of the OT signaling pathway, such as OT, the OT receptor (OTR), and CD38, consistently support its critical role in social behaviors (21, 25, 26). When abundant OT release occurs in the brain, for instance during lactation, AI is one of the most robustly activated brain regions (27, 28). Nevertheless, quantitative analyses for the distribution of OT fibers in multiple brain regions have reported that the OT projections to the AI are negligible in multiple species (23, 29). In contrast, OT projection and rich expression of OTR binding are found in the PI (23, 30). Thus, the oxytocin signaling may be distinct between the PI and AI. Together, the mechanism that underlies the activation of the AI in response to OT in humans remains to be elucidated.

We have recently proposed the importance of using animal models in research to discover novel translatable mechanisms at the molecular and cellular level in this understudied brain region (18). Outstanding studies with rodents that explore roles of the insular cortex, in particular focusing on the PI, have indeed emerged in the past several years: at least in rodents, the PI is involved in emotional processing and resultant social behavior in response to socially affective stimuli is mediated by the OTR-dependent neural activity (31, 32). In contrast, we addressed the aforementioned enigma from human imaging study and decipher a role of AI in social behavior in association with OT signaling. We used an AI-enriched molecule Cyp26B1, which is crucial for retinoic acid (RA) degradation and involved in the pathology of neuropsychiatric conditions, as a molecular lead to decipher the role of AI in social cognition.

## Methods

For experimental methods, please see the supplementary method section.

### Statistical analysis

Data were analyzed using the statistical methods stated in each figure legend, and all statistical analyses were performed using SPSS. Sample sizes were chosen on the basis of previous studies. When performing parametric tests such as t-tests and ANOVAs, the assumptions of normality (normal distribution) and homogeneity (similar variance among groups) were tested using Shapiro-Wilk’s test and Levene’s test, respectively. Data were mostly analyzed by one-way ANOVAs, two-way ANOVAs, and two-way repeated measures ANOVAs followed by post hoc test. One-way ANOVAs followed by Tukey’s multiple comparisons were performed for testing statistical difference between means of three or more independent groups. Two-way ANOVAs followed by Bonferroni post hoc tests were used when examining the influence of two different factors on a dependent variable. However, if any repeated factor was present, two-way repeated measures ANOVAs followed by Bonferroni post hoc tests were used. For simple pair-wise comparisons, Student’s *t*-tests were used.

## Results

### The AI underlies social novelty recognition

We first investigated whether the AI is a critical brain area for social cognition in mice. We introduced bilateral excitotoxic lesions in either the AI or its adjacent brain region, the orbitofrontal cortex (OFC), in C57BL6/J wild-type male mice (see **Figure S1** in the online supplement). In 3-chamber social interaction test, normal mice prefer the enclosure with a social subject (mouse) (trial 1: sociability) and show a selective preference to a novel mouse compared with a familiar mouse (trial 2: social novelty recognition) (**Figure 1A**). We observed a deficit only in trial 2 in AI-lesioned mice, whereas OFC-lesioned mice normally behaved in both trials (**Figure 1A**). Thus, AI-lesioned mice, but not OFC-lesioned mice, exhibited a significant deficit in social novelty recognition but not in sociability in this assay. To address whether the deficit was specific to the social context, we employed a 5-trial social novelty recognition paradigm guided by social or non-social cues. Both OFC-lesioned and AI-lesioned mice displayed normal behaviors for non-social cue (odor): gradually reducing exploration time to a banana-scent during 4 consecutive trials, but substantially increasing exploration when the scent was changed to a strawberry-scent on the fifth trial (**Figure 1B**). In the context of social cue, the OFC-lesioned mice also displayed a normal response (**Figure 1C**). In contrast, AI-lesioned mice showed significant deficits in response to social cue: absence of a habituation during the first 4 trials and no increase in exploration when exposed to a new conspecific cue in the fifth trial (**Figure 1C**). To avoid the possibility that these AI-specific deficits are associated with olfactory function or locomotor activity, we examined the hidden food test and open field test and showed that these are not confounding factors (see **Figure S2A** and **S2B** in the online supplement). In addition, AI-lesioned female mice behaved similarly to male mice, displaying a deficit in social novelty recognition (see **Figure S2C** in the online supplement). Together, these results indicate that the AI plays a role in social novelty recognition in mice, which clearly differentiates it from the OFC.

**FIGURE 1.**
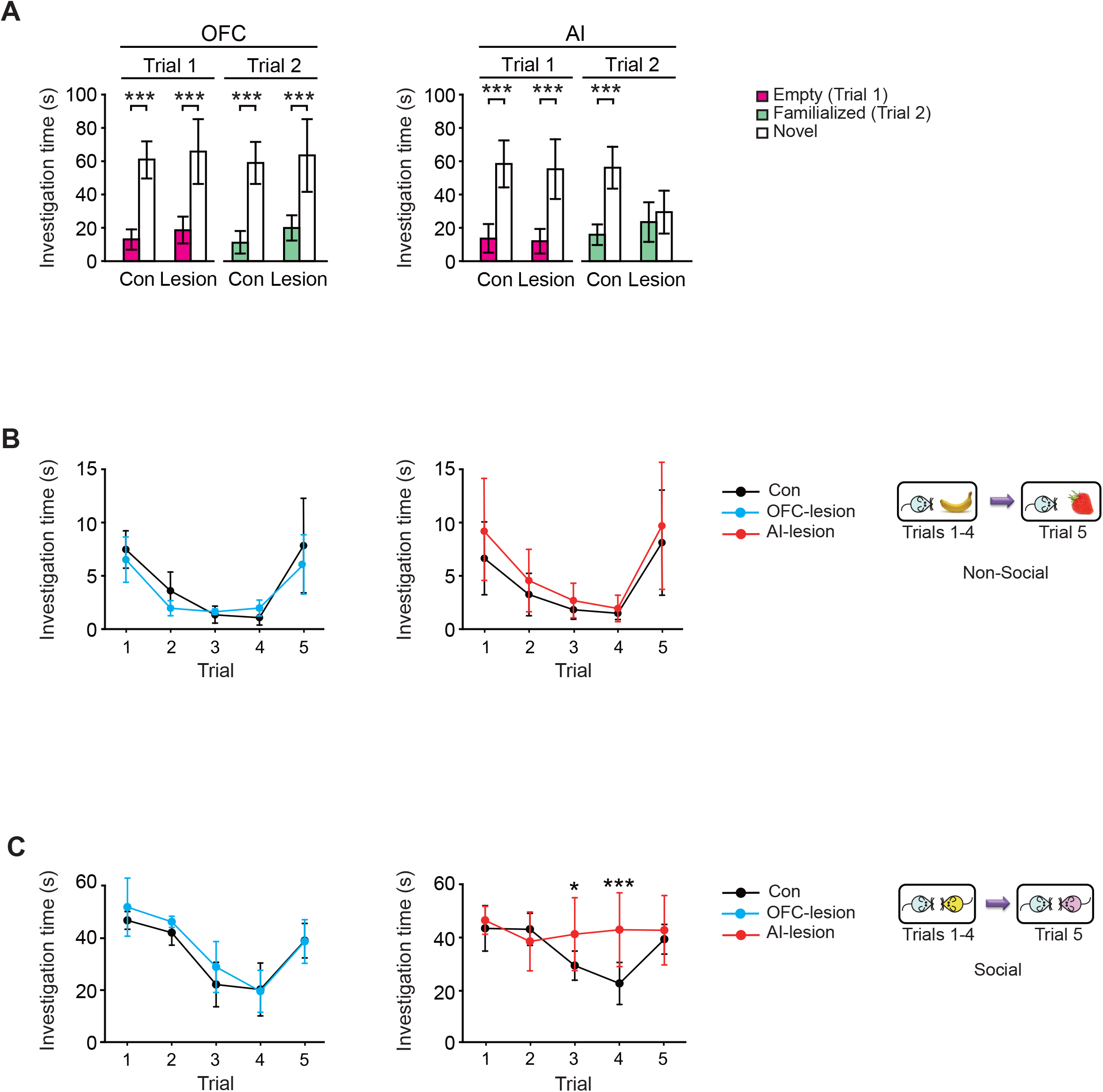
The AI underlies social novelty recognition. **Panel A**, Left: OFC-lesioned mice show normal performance in both sociability (Trial 1) and social novelty recognition (Trial 2) in the three-chamber social interaction test. Right: AI-lesioned mice show a deficit in social novelty recognition [Trial 2: AI-Con, *P* < 0.001 and AI-lesion, *P* = 0.103; multiplicity-adjusted *P* values; *F*_lesion x chamber_ (1, 72) = 44.72, *P* < 0.001] but exhibited normal sociability (Trial 1). OFC: control (Con), n = 12; lesion, n = 15. AI: Con, n = 19; lesion, n = 19. **Panel B**, Both OFC-lesioned (left) and AI-lesioned (right) mice exhibit normal performance in the 5-trial recognition test with non-social cues. OFC-Con, n = 5; OFC-lesion, n = 4; AI-Con, n = 8; and AI-lesion, n = 8. **Panel C**, OFC-lesioned mice exhibit normal performance in the 5-trial recognition test with social cues (left), while AI-lesioned mice exhibit abnormal performance in the 5-trial recognition test with social cues (right). AI-lesioned mice are significantly differed from AI-Con [*F*_lesion x trial_ (4, 56) = 4.705, *P* < 0.01]. OFC-Con, n = 5; OFC-lesion, n = 4; AI-Con, n = 8; and AI-lesion, n = 8. Two-way ANOVA with Bonferroni *post-hoc* test for **A**, and two-way repeated measures ANOVA with Bonferroni *post-hoc* test for **B** and **C**. Data are represented as mean ± S.D. *\**p<0.05, *\*\** p<0.005, *\*\*\** p<0.001.

### Alteration of RA signaling in the AI leads to abnormal social novelty recognition

To study the molecular, cellular, and circuitry mechanism that underlies this AI-mediated behavior, we looked for a lead molecule to build a useful mouse model at both the basic and translational science levels. We hypothesized that the target would be enriched in the layer 5 pyramidal neurons of the AI, whose gene is significantly highlighted in the genome-wide association studies (GWAS) for brain disorders. We combined the information from our own microarray data of fluorescence-activated cell sorting (FACS)-isolated AI layer 5 pyramidal neurons (see Methods for details) and an anatomic gene expression atlas (AGEA) (33), which were cross-referenced with the GWAS risk genes so that we highlighted 47 candidate molecules (see **Figure S3A** in the online supplement). Following successful precedents in which mouse models that genetically modify enzymes critical to the biosynthesis of key mediators (e.g., serine racemase for D-serine and nitric oxide synthase for nitric oxide) have been highly useful in exploring neurobiological mechanisms (34, 35), we further narrowed the candidates down to two molecules that fall in this category, Cyp26B1 [cytochrome P450 26B1: a retinoic acid (RA)-degrading enzyme] (36, 37) and Ckb (creatine kinase brain type) (see **Figure S3A** in the online supplement). Cyp26B1 has been highlighted by an analysis using a comprehensive transcriptome study of postmortem brains from patients with autism spectrum disorder, schizophrenia and bipolar disorder (see **Figure S3B** in the online supplement) (38). Additionally, RA signaling is one of the top 4 signaling cascades in brain disorders (**Figure 2A**). Furthermore, we observed that a chronic infusion of RA locally into the AI resulted in a phenocopy of the AI-lesion, exhibiting a significant deficit in social novelty recognition (trial 2) but not in sociability (trial 1) in the three-chamber social interaction test (**Figure 2B** and see **Figure S4A** in the online supplement). Taken together, we decided to study the unique role of the AI in social novelty recognition by using RA signaling, in particular Cyp26B1, as a molecular lead.

**FIGURE 2.**
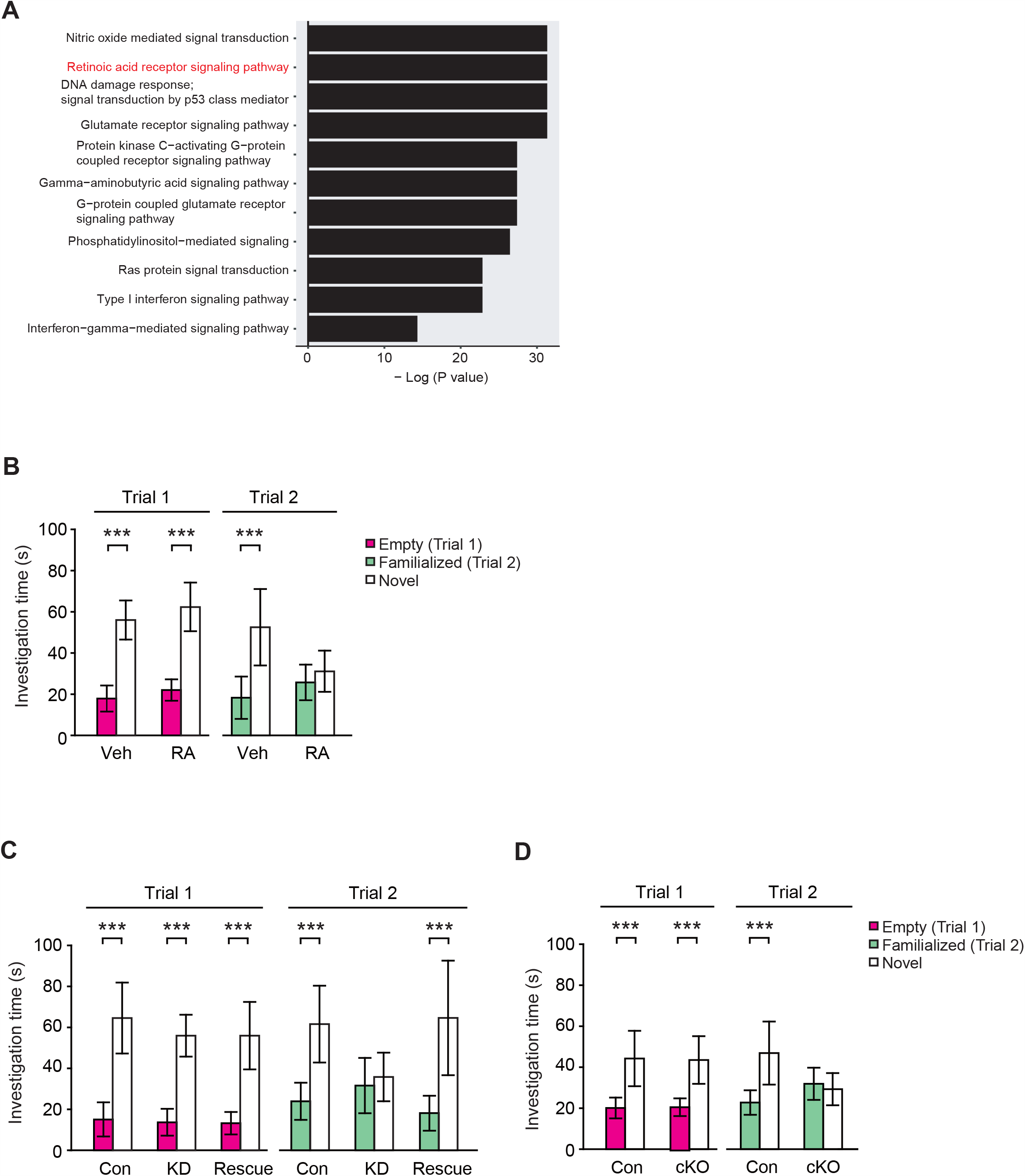
RA signaling in the AI underlies social novelty recognition. **Panel A**, Signaling pathways critical to brain disorders. RA signaling is underscored as one of the top 4. **Panel B**, Chronic intra-insular administration of RA leads to a deficit in social novelty recognition [Veh, *P* < 0.001 and RA, *P* = 0.467; *F*_treat x chamber_ (1, 20) = 8.014, *P* < 0.05], but not in sociability. Vehicle (Veh), n = 6; and RA, n = 6. **Panel C**, Reduced Cyp26B1 expression in the AI by RNAi-mediated knockdown (KD) causes a significant deficit [Con, *P* < 0.001, KD, *P* = 0.502, and rescue, *P* < 0.001; *F*_AAV x chamber_ (2, 60) = 11.38, *P* < 0.001] in social novelty recognition, but not in sociability. Con, n = 11; KD, n = 13; and KD + Cyp26B1^R^ (Rescue), n = 9. **Panel D**, Cyp26B1 conditional knockout specifically in the AI (cKO) phenocopies [Con, *P* < 0.001 and cKO, *P* = 0.545; *F*_AAV x chamber_ (1, 42) = 18.59, *P* < 0.001] the deficits of social novelty recognition that were found in the AI-Cyp26B1 KD mice. AAV-αCaMKII-EYFP (Con), n = 11; and AAV-αCaMKII-GFP-Cre (cKO), n = 12. Two-way ANOVA with Bonferroni *post-hoc* test for **B, C**, and **D**. Data are represented as mean ± S.D. *\**p<0.05, *\*\** p<0.005, *\*\*\** p<0.001.

To directly examine the role of Cyp26B1 in the AI on social behavior, we bilaterally knocked down the expression of Cyp26B1 in the AI by adeno-associated virus (AAV)-mediated RNAi [AAV-U6-shCyp26B1-CMV-mCherrry (AAV-Cyp26B1 KD)] (see **Figure S4B** in the online supplement). Efficient knockdown (KD) of Cyp26B1 and a reciprocal augmentation of RA signaling were confirmed *in vivo* (see **Figure S5A** and **S5B** in the online supplement). Consistent with the effect of local RA infusion into the AI (**Figure 2B**), Cyp26B1 KD in the AI (AI-Cyp26B1 KD) strongly induced a deficit in social novelty recognition without affecting sociability in the three-chamber social interaction test whereas control mice (AI-Con) injected with AAV-U6-shControl-CMV-mCherrry (AAV-Con) displayed typical social behaviors (**Figure 2C**). Restoring the expression of Cyp26B1 by introducing RNAi-resistant Cyp26B1 (Cyp26B1^R^) with AAV-hSyn-Cyp26B1^R^-HA (AAV-Cyp26B1^R^) in the AI rescued the deficit in social novelty recognition elicited by Cyp26B1 loss-of-function (**Figure 2C**). The AI-Cyp26B1 KD mice also displayed recognition deficits in the context of social but not non-social cues (see **Figure S5C** in the online supplement). These data indicate the direct involvement of Cyp26B1 in the AI in social novelty recognition. To confirm and extend the results obtained from the RNAi approach, we used Cyp26B1 conditional knockout mice (39): we selectively depleted Cyp26B1 in αCaMKII-positive cells in the AI by injecting AAV-αCaMKII::GFP-Cre in the AI of *Cyp26B1*-floxed (*Cyp26B1*^*flox/flox*^) mice (see **Figure S4C** in the online supplement). The genetic depletion of Cyp26B1 in αCaMKII-positive cells (pyramidal neurons) of the AI led to deficits in social novelty recognition (**Figure 2D**), which is consistent with the phenotype in AI-Cyp26B1 KD mice.

### AI-mediated social novelty recognition is maintained by proper activity of the layer 5 pyramidal neurons for which RA-mediated gene transcription plays a role

Because RA signaling is primarily involved in transcriptional regulation, we hypothesized that Cyp26B1 KD/KO affected these behavioral phenotypes via altered gene transcription. Thus, we performed an unbiased molecular profiling analysis of FACS-enriched AI cortical layer 5 pyramidal neurons. There was a small subset of differentially expressed genes between control and KD groups [false discovery rate (FDR) < 0.1], while the expression profiles of control and rescue groups were similar with good reproducibility among biological triplicates (**Figure 3A**). The majority of the genes we identified (15 out of 20) included a retinoic acid response element (RARE) (40) and/or were known to bind with retinoic acid receptors (RARs) (41) (see **Table S1** in the online supplement), suggesting that these genes are likely to be regulated by RA signaling. Results from quantitative real-time PCR analyses were consistent with the molecular profiling data (see **Figure S5D** in the online supplement). Bioinformatic analyses strongly suggested that chronically upregulated RA signaling might cause abnormal dendritic spine regulation in the AI layer 5 pyramidal neurons (**Figure 3B**). Consistent with this prediction, we experimentally confirmed that AI-Cyp26B1 KD caused a significant reduction in spine density in the proximal apical dendrites, compared to the control group, while the expression of Cyp26B1^R^ effectively normalized this deficit (**Figure 3C**). These results suggest that the decreased expression of Cyp26B1 was the sole cause for the spine deficit.

**FIGURE 3.**
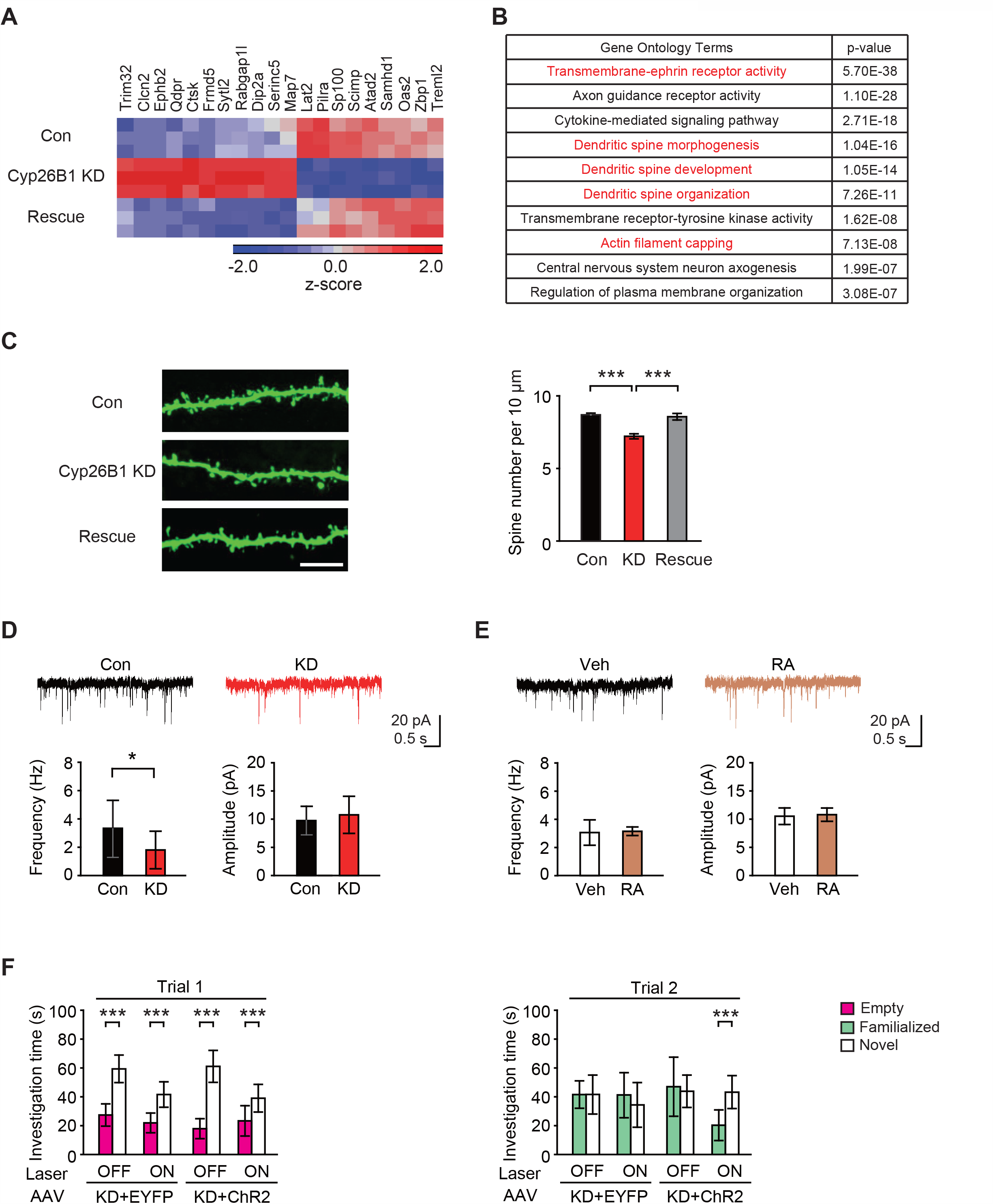
AI-associated social novelty recognition is mediated by proper activity of the AI layer 5 pyramidal neurons. **Panel A**, A heat map for differentially expressed genes from a molecular profiling analysis of FACS-enriched AI layer 5 pyramidal neurons. Three brains per group were used for sampling after injection of AAV-Con (Con), AAV-Cyp26B1 KD (Cyp26B1 KD), and AAV-Cyp26B1 KD + AAV-Cyp26B1^R^ (Rescue), respectively. Expression levels are normalized using z-score transformation and represented as a color scale of the z-score. **Panel B**, The top 10 terms of gene ontology enrichment analyses. Groups associated with dendritic spine plasticity are highlighted in red. **Panel C**, Spine densities of the YFP-expressing layer 5 pyramidal neurons in the AI of Thy1-YFP-H transgenic mice are reduced by Cyp26B1 KD [*F*(2, 41) = 20.67, *P* < 0.001]. Left: representative fluorescence images of the proximal apical dendritic spines. Scale bar, 10 μm. Right: quantitative assessment of averaged spine densities from 3–4 brains per group. Con, n = 13; KD, n = 16; and Rescue, n = 15 neurons. **Panel D**, The AI layer 5 pyramidal neurons infected with AAV-Cyp26B1 KD (KD) show a decrease in sEPSC frequency [left: *t*(19) = 0.217, *P* < 0.05], but not amplitude (right), compared to neurons infected with AAV-Con (Con). Top: representative sEPSCs traces recorded from AAV-infected neurons. Con, n = 10 neurons from 5 mice; and KD, n = 11 neurons from 4 mice. **Panel E**, Acute RA treatment does not change the frequency (left) or amplitude (right) of sEPSCs in the AI layer 5 pyramidal neurons compared to vehicle-treated cells (Veh) from wild-type mice. Top: representative sEPSCs traces recorded from neurons in brain slices treated with Veh or RA. n = 10 neurons from 5 mice per group. **Panel F**, Optogenetic excitation of the AI pyramidal neurons during the three-chamber social interaction test does not affect sociability (trial 1) in AI-Cyp26B1 KD mice. Optogenetic excitation restores proper social novelty recognition that was deficient in AI-Cyp26B1 KD mice [KD + ChR2: laser-OFF, *P* = 0.615 and laser-ON, *P* < 0.001; *F*_laser x chamber_ (1, 36) = 8.653, *P* < 0.01]. KD + EYFP, n = 8; and KD + ChR2, n = 10. One-way ANOVA with *post-hoc* Tukey’s multiple comparison test for **C**, two-tailed *t* test for **D**, and **E**, two-way repeated measures ANOVA with Bonferroni *post-hoc* test for **F**. Data are represented as mean ± S.D. *\**p<0.05, *\*\** p<0.005, *\*\*\** p<0.001.

We also validated the spine changes at the electrophysiological level. The AI layer 5 pyramidal neurons in acute slices from AI-Cyp26B1 KD mice exhibited a significantly lower frequency of spontaneous excitatory postsynaptic currents (sEPSCs) compared to those from control mice, whereas both groups of neurons showed no difference in amplitude (**Figure 3D**). The decrease in sEPSC frequency in the AI layer 5 pyramidal neurons implies that there would be reduced activity in these neurons and their downstream targets. In contrast, acute bath application of RA to brain slices from wild-type mice did not affect the sEPSCs of the AI layer 5 pyramidal neurons (**Figure 3E**). Taken together, we showed that proper control of RA signaling is critically required for the maintenance of dendritic architecture and activity of the AI layer 5 pyramidal neurons. Thus, we optogenetically activated the AI pyramidal neurons during the three-chamber social interaction test in mice infected with Cyp26B1 KD AAV and channel rhodopsin (ChR2)-EYFP AAV (see **Figure S6** in the online supplement). Optogenetic excitation of the AI pyramidal neurons did not affect sociability (trial 1) in the AI-Cyp26B1 KD mice (**Figure 3F**).

In contrast, optogenetic excitation in the AI restored proper social novelty recognition (trial 2) (**Figure 3F**). This data suggests that modulation of spontaneous neuronal activity of the AI layer 5 pyramidal neurons could affect social novelty recognition.

### OT influences AI-mediated social novelty recognition

Next, we looked for circuitry mechanisms by which the activity of AI pyramidal neurons is properly maintained in social contexts. We examined plausible afferent projections to the AI by using a retrograde tracing method in which we injected cholera toxin subunit B-conjugated with Alexa 488 (CTB-Alexa 488) into the AI (**Figure 4A**). As expected, we observed a strong signal from the contralateral AI and basolateral amygdala, well-known brain regions that tightly connect with the AI (**Figure 4B**) (42, 43). Then we assessed possible projections to the AI from the hypothalamic paraventricular nucleus (PVN) and supraoptic nucleus (SON), where OT neurons originate (44). A recent publication reported that OT could directly influence social affective behavior mediated by the posterior insula (PI) (32). Nevertheless, it remains elusive how OT affects the function of the AI. It is well known that the AI and PI (see **Figure S1A** in the online supplement) are quite distinct anatomically and functionally in many aspects (45, 46).

**FIGURE 4.**
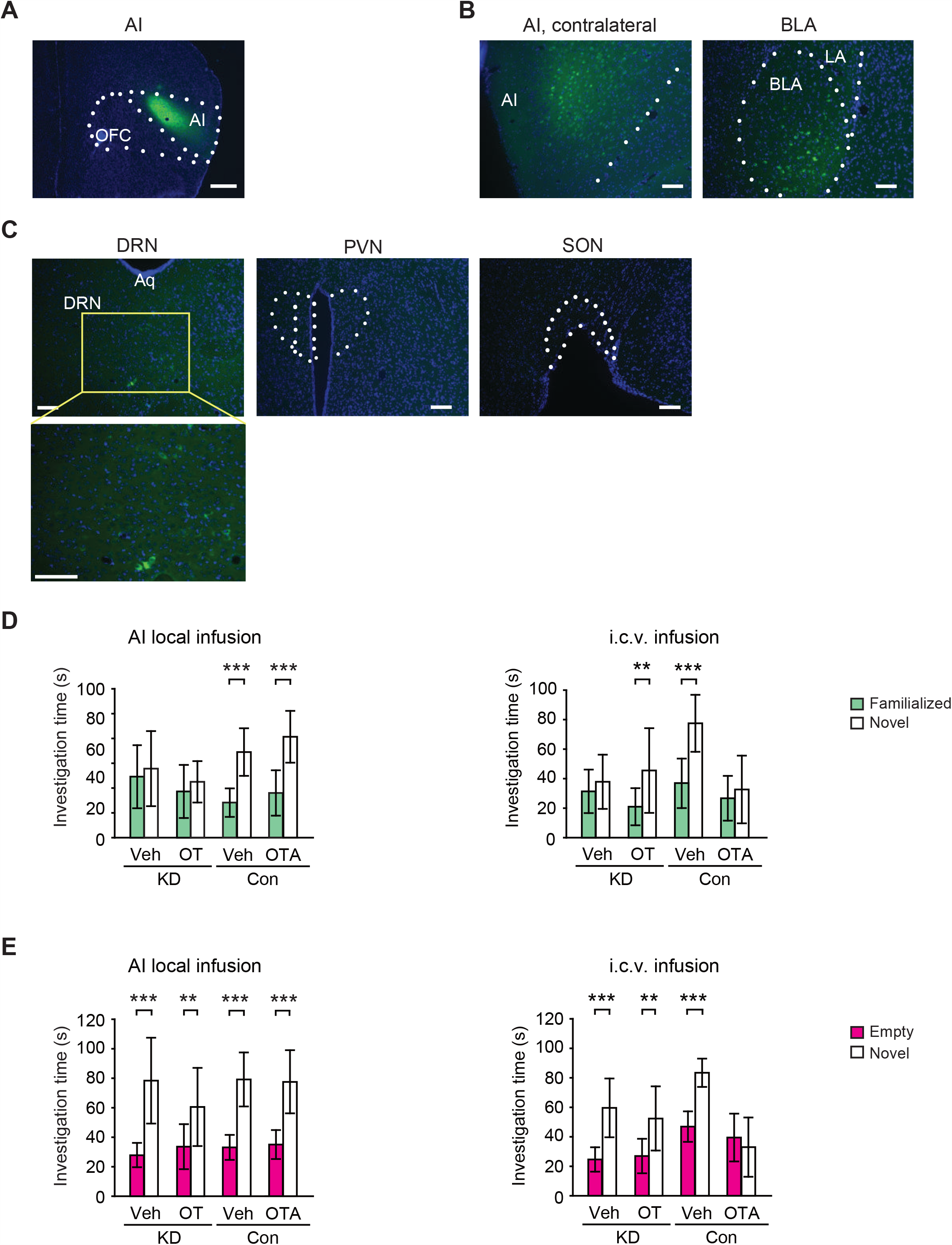
AI-mediated social novelty recognition is regulated by OT in a non-classic mechanism. **Panel A**, Representative image for the unilateral injection of green fluorescence retrograde tracer (CTB-Alexa 488) into the right AI. Scale bar, 500 μm. **Panel B, C**, Representative images showing retrograde labeled green fluorescence-positive cell bodies in AI-projecting brain areas. The two brain regions [contralateral AI and basolateral amygdala (BLA)] known to have afferent projections to the AI are displayed (**B**) as positive controls, together with other brain regions of interest (**C**). Among the DRN, PVN and SON, green fluorescence retrogradely transported from the AI was only observed in the DRN. LA, lateral amygdala. Aq, aqueduct. Scale bars: 100 μm. **Panel D**, Effect of OT and OTA on social novelty recognition. Left: local infusion of OT or OTA into the bilateral AI show no effect on AI-Cyp26B1 KD (KD) or control (Con) mice, respectively. KD-Veh, n = 13; KD-OT, n = 11; Con-Veh, n = 11; and Con-OTA, n = 12. Right: i.c.v. administration of OT normalizes the social novelty recognition deficit in AI-Cyp26B1 KD mice (KD) [Veh, *P* = 0.463 and OT, *P* < 0.01; *F*_drug x chamber_ (1, 36) = 2.122, *P* = 0.154], while OTA disrupts proper social novelty recognition in healthy control mice (Con) [Veh, *P* < 0.001 and OTA, *P* = 0.507; *F*_drug x chamber_ (1, 32) = 7.627, *P* < 0.01]. KD-Veh, n = 10; KD-OT, n = 10; Con-Veh, n = 9; and Con-OTA, n = 9. **Panel E**, Effect of OT and OTA on sociability. Left: local infusion of OT or OTA into the bilateral AI show no effect on sociability in both AI-Cyp26B1 KD and control mice. KD-Veh, n = 13; KD-OT, n = 11; Con-Veh, n = 11; and Con-OTA, n = 12. Right: i.c.v. administration of OT has no effect on sociability in the AI-Cyp26B1 KD mice, while OTA disrupts sociability in control mice [Veh, *P* < 0.001 and OTA, *P* = 0.355; *F*_drug x chamber_ (1, 32) = 19.28, *P* < 0.001]. KD-Veh, n = 10; KD-OT, n = 10; Con-Veh, n = 9; and Con-OTA, n = 9. Two-way ANOVA with Bonferroni *post-hoc* test for **D** and **E**. Data are represented as mean ± S.D. *\*P* < 0.05, *\*\*P* < 0.005, *\*\*\*P* < 0.001.

Several reports have indicated that the projection of OT neurons to the AI is quite sparse or negligible (23, 29). Our experimental data were indeed in accordance with these reports, indicating that there are no detectable projections to the AI from the hypothalamic PVN and SON (**Figure 4C**).

If OT does not influence the AI neurons via direct projection of OT neurons, another scenario is via diffusion of OT into the AI. To test this possibility, we locally infused OT into the AI of AI-Cyp26B1 KD mice and examined whether exogenous OT could ameliorate the AI-associated social novelty recognition deficits (see **Figure S7A** in the online supplement). Local infusion of OT into the AI of AI-Cyp26B1 KD mice did not alter their deficit in social novelty recognition (**Figure 4D**). We also locally infused desGly-NH_2_-d(CH_2_)_5_ [D-Tyr^2^, Thr^4^] OVT, a selective OTR antagonist (OTA), into the AI of control mice and did not observe any changes in social behavior (**Figure 4D** and see **Figure S7A** in the online supplement). Nevertheless, intracerebroventricular (i.c.v.) administration of OT normalized the social novelty recognition deficit in AI-Cyp26B1 KD mice in which RA-signaling is selectively impaired only in the AI (**Figure 4D** and see **Figure S7B** in the online supplement). The OTA also influenced social novelty recognition in control mice by the i.c.v. administration (**Figure 4D** and see **Figure S7B** in the online supplement). These effects were selective to social novelty recognition, except i.c.v. administration of OTA that also altered sociability (**Figure 4E**). Together, these results suggest that, although OT specifically participates in the regulation of the AI-mediated social novelty recognition, this is unlikely to be via the influence of OT directly on the AI neurons. Instead, OT may influence AI-mediated social novelty recognition indirectly, by remotely affecting the neurons that project to the AI from different brain regions.

### Serotonin influences AI-associated social novelty recognition via 5-HT2C receptor

In the retrograde tracing experiments to look for neuronal projections to the AI, we examined several brain regions in addition to the PVN and SON. Among them, we observed a distinct signal in the dorsal raphe nucleus (DRN) (**Figure 4C**), which is known to provide major serotonergic projections to the forebrain. Thus, we examined how modulation of serotonergic inputs might affect AI-mediated social novelty recognition. Based on their high expression levels in the AI (http://mouse.brain-map.org) we focused on two serotonin receptors (5-HT1A and 5-HT2C). First we tested the effects of their agonists and antagonists by systemic injection: only CP 809101 (a 5-HT2C receptor agonist) normalized the social novelty recognition deficit in AI-Cyp26B1 KD mice, whereas SB 242084 (a 5-HT2C receptor antagonist) disrupted social novelty recognition in control mice (see **Figure S8A** in the online supplement). To further validate that the impact of the agonist and antagonist on behavior is directly mediated via the AI, we conducted local infusion experiments. Stimulating the AI serotonergic system via 5-HT2C normalized the social novelty recognition deficit in AI-Cyp26B1 KD mice (**Figure 5A**), while inhibition of 5-HT2C in the AI of control mice impaired social novelty recognition (**Figure 5A**). Treatment with the 5-HT2C receptor agonist increased both the amplitude and frequency of sEPSCs in AI-Cyp26B1 KD pyramidal neurons without affecting presynaptic features (**Figure 5B** and see **Figure S8B** in the online supplement). These data suggest that the 5-HT2C receptor agonist ameliorated deficits in the AI layer 5 pyramidal neurons of AI-Cyp26B1 KD mice.

**FIGURE 5.**
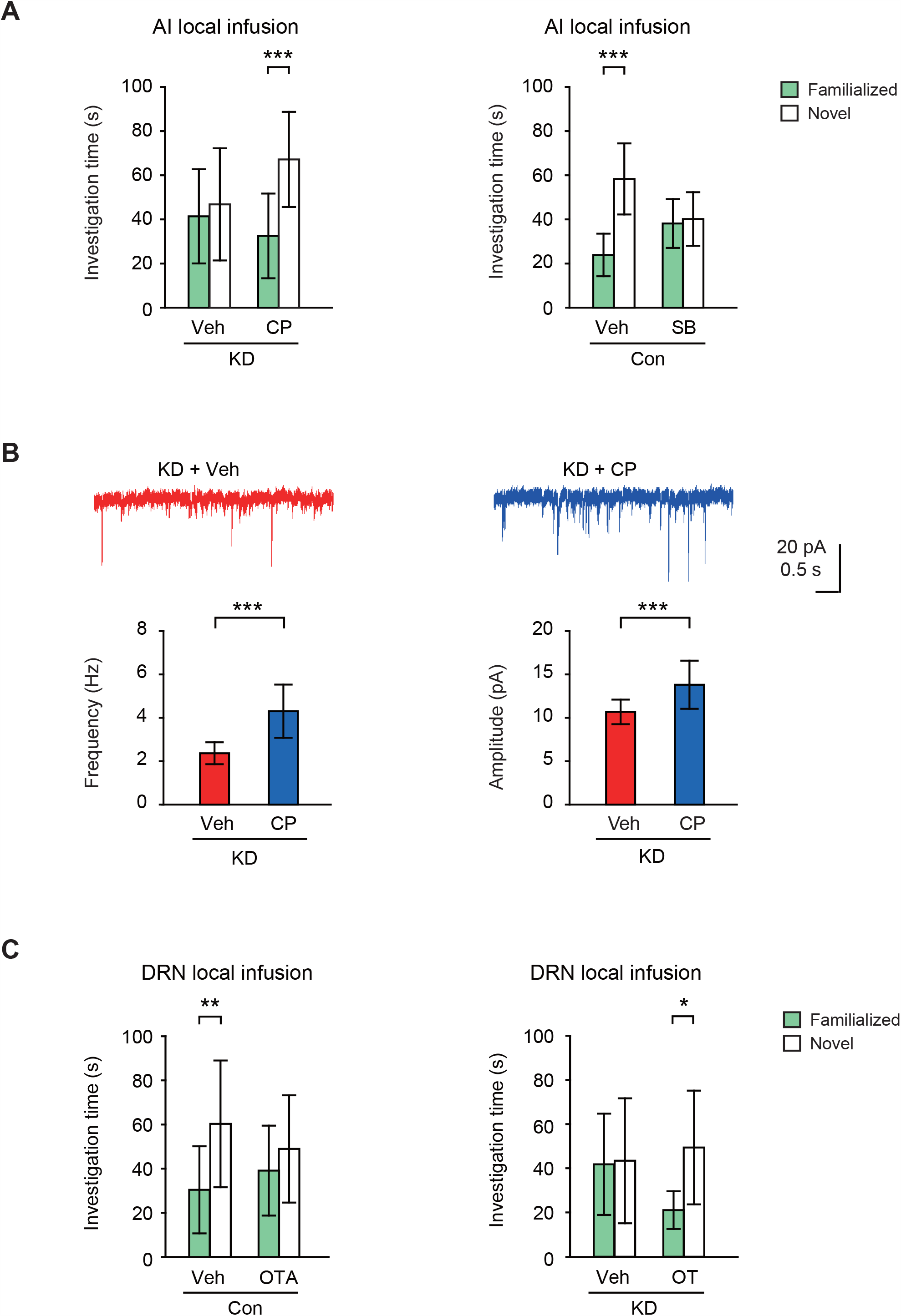
AI-mediated social novelty recognition is regulated by serotonin via the 5-HT2C receptor expressed in the AI. **Panel A**, The AI 5-HT2C receptor as an intervention for altered social novelty recognition. Left: bilateral local infusion of the 5-HT2C receptor agonist CP 809101 (CP) into the AI normalizes the altered social novelty recognition in AI-Cyp26B1 KD mice (KD) [Veh, *P* = 0.534 and CP, *P* < 0.001; *F*_drug x chamber_ (1, 48) = 5.738, *P* < 0.05]. KD-Veh, n = 13; and KD-CP, n = 13. Right: bilateral local infusion of the 5-HT2C receptor antagonist SB 242084 (SB) into the AI disrupts social novelty recognition in control mice (Con) [Veh, *P* < 0.001 and SB, *P* = 0.718; *F*_drug x chamber_ (1, 38) = 17.51, *P* < 0.001]. Con-Veh, n = 11; and Con-SB, n = 11. **Panel B**, Acute activation of 5-HT2C receptors increases both the frequency [left: *t*(27) = 5.659, *P* < 0.001] and amplitude [right: *t*(27) = 3.955, *P* < 0.001] of sEPSCs in the AI layer 5 pyramidal neurons from AI-Cyp26B1 KD mice. KD + Veh, n = 14 neurons from 5 mice; and KD + CP, n = 15 neurons from 4 mice. **Panel C**, OT and OTA in the DRN modulates AI-mediated social novelty recognition. Left: local infusion of OTA into the DRN disrupts normal social novelty recognition in control mice (Con) [Veh, *P* < 0.01 and OTA, *P* = 0.385; *F*_drug x chamber_ (1, 40) = 1.956, *P* = 0.170]. Con-Veh, n = 12; and Con-OTA, n = 10. Right: local infusion of OT into the DRN restores proper social novelty recognition in AI-Cyp26B1 KD mice [Veh, *P* = 0.881 and OT, *P* < 0.05; *F*_drug x chamber_ (1, 30) = 2.901, *P* = 0.099]. KD-Veh, n = 9; and KD-OT, n = 8. Two-way ANOVA with Bonferroni *post-hoc* test for **A** and **C**. Two-tailed *t*-test for **B**. Data are represented as mean ± S.D. *\*P* < 0.05, *\*\*P* < 0.005, *\*\*\*P* < 0.001.

### Modulation of DRN neurons by OT influences AI-associated social novelty recognition

We have shown that AI-mediated social novelty recognition is properly maintained by serotonergic regulation via 5-HT2C receptors expressed on AI pyramidal neurons. Meanwhile, the same social behavior is also regulated by OT, which likely plays a role in the AI by remotely affecting the neurons that project to the AI from different brain regions. Given that serotonergic innervation in the AI is from the DRN, one possible hypothesis is that OT directly influences the neurons in the DRN, which in turn regulates the activity of the AI pyramidal neurons. Indeed, OT neurons from the PVN reportedly project to the DRN (24), where OT receptors are reportedly highly expressed (47). To test our hypothesis, first we infused OTA specifically into the DRN of control mice and tested its effects on social novelty recognition. We observed that this treatment selectively impaired social novelty recognition (**Figure 5C** and see **Figure S9** in the online supplement). To test whether interfering with OT signaling in the DRN is specifically associated with the AI-mediated social novelty recognition, we infused OT specifically into the DRN of AI-Cyp26B1 KD mice. OT infusion into the DRN significantly restored social novelty recognition (**Figure 5C** and see **Figure S9** in the online supplement).

## Discussion

In the present study, by using mouse models, we report a pivotal role of the AI in social novelty recognition in which proper activity of the layer 5 pyramidal neurons is required. At the molecular and cellular level, the AI-mediated social novelty recognition is maintained by proper activity of AI layer 5 pyramidal neurons, for which retinoic acid-mediated gene transcription plays a role. Social novelty recognition is also regulated by 5-HT2C receptor expressed in the AI. Furthermore, we demonstrate that OT influences the AI-mediated social cognition not by direct projection of OT neurons, nor by direct diffusion of OT in the AI, but rather affecting OTR-expressing neurons in the DRN where serotonergic neurons are projected to the AI. Accordingly, the regulatory mechanism by OT on the AI is distinct from that on the PI where direct projection of OT neurons plays an important role.

In the brain, OT exerts its control through long-range axonal release, hence projection of OT fibers and the expression of OTR in the target brain region is crucial. In this context, several brain regions have been postulated to be involved in OT-associated social cognition, including prefrontal cortex, hippocampus, and olfactory nucleus (48–51), and more recently the PI (32). Unlike the PI, projection of OT neurons to the AI is sparse (32) and the expression of OTR is attenuated (52). In human functional connectivity study, there are distinct subdivisions in the insula cortex for processing cognitive information and affective feeling (43). Recent studies in rodent models have highlighted a role of the PI in social behavior (31, 32). It is an important future question whether and how the involvement of the AI and PI in social cognition and behavior is similar and different, possibly with distinct regulatory mechanisms for each sub-region, including those by OT.

We underscored the significance of the 5HT2C receptor expressed in the AI for social novelty recognition. In the past, through systemic administration of a 5-HT2C receptor antagonist in wild-type animals (53) or the phenotype of 5-HT2C receptor genetic knockout (54), the involvement of this receptor in both social novelty recognition and sociability has been suggested. In contrast to a broad expression of 5-HT1A and 2A receptors in the frontal cortex (55, 56), the expression of 5-HT2C receptor is more restricted with a more prominent expression in the medical prefrontal cortex and AI (56, 57). Our data in the present study showed that local injection of a 5-HT2C receptor antagonist elicits the deficits only in social novelty recognition, but not in sociability, which is compatible to our mechanistic proposal. Thus, the impact of 5-HT2C receptor in sociability may be via the receptor in the medial prefrontal cortex that is known to be involved in sociability (58, 59). Genetic manipulations of this receptor in the AI pyramidal neurons further can validate its specific role in social novelty recognition.

The present study has pinned down the mechanistic question for the next step of how OT can influence the AI-mediated social novelty recognition in a non-classic (neither OT neuron direct projection, nor direct diffusion of OT to the AI) way. One of the most tempting working hypotheses may be a disynaptic model in which the OTR-expressing DRN serotonin neurons, receiving input from PVN/SON-originated OT neurons, directly project to the AI pyramidal neurons and exert their effect on the AI-dependent social novelty recognition. This model is theoretically addressable, although experimental optimization may be very difficult, by depleting OTR specifically from the AI-projecting DRN neurons and further rescue the AI projection by activating this projection specifically using optogenetic manipulation. Through this experiment, the necessity and sufficiency of the disynaptic mechanism may be shown in future studies to account for the influence of OT on the AI-mediated social novelty recognition. Alternately, AI may be influenced by OT through an indirect mechanism via a brain area to which OTR-expressing non-serotonergic neurons project from the DRN.

Future investigation may also address local mechanisms in greater extent, for instance how AI pyramidal neuron activity is regulated for AI-mediated social novelty recognition. These mechanisms may include modulation of neuronal excitability possibly mediated via G-protein coupled potassium current downstream of 5HT2C receptor (60, 61), and local synaptic regulation via RA signaling. Future studies should also address downstream executive mechanisms whereby the AI pyramidal neurons eventually influence behavior. The functional relationship of the AI with the amygdala and hippocampus, in particular the CA2 (62) and ventral CA1 (63), may be topics of future investigations.

Although recent human genetic research has suggested the significance of RA-associated genes in brain disorders (64, 65), the biological role of the RA-associated cascade in mental conditions has not yet been fully addressed. Mild elevation of RA signaling by RARβ2 expression or by chronic neuronal inactivation have been reported to show a positive effect on neurite regeneration and homeostatic synaptic scaling (66–69). In contrast, in our experimental mouse model, we chronically disturbed RA signaling by suppressing Cyp26B1 in the AI layer 5 pyramidal neurons. The present model will be useful to decipher further mechanisms of how OT and serotonin regulate AI-mediated social cognition. Furthermore, the model will also be useful to study pathological impact of chronic disturbance in the RA signaling in the AI. Consistent with this notion, in multiple expression studies of postmortem brains from patients with mental disorder (38), Cyp26B1 has been underscored as a top molecule of interest. Together, the Cyp26B1 models used in the present study will be useful to explore mechanisms of social cognition deficits in mental disorders.

## Acknowledgments

We thank Yukiko Y. Lema for organizing the manuscript and Dr. Melissa Landek-Salgado for critical reading and discussion of the manuscript. We also thank Lauren Guttman for reading the manuscript and her comments. We thank Dr. Maria I. Morasso (NIAMS, NIH) and Dr. Hiroshi Hamada (RIKEN) for mice carrying the floxed *Cyp26B1* allele (*Cyp26b1*^*f/f*^). The human synapsin1 (hSyn) promoter-containing AAV plasmid (AAV-6P-SEWB) was a gift from Dr.

Sebastian Kügler (Univ. of Göttingen). This work was supported by NIH grants MH-094268 Silvio O. Conte center and MH-105660 (A.S.), as well as foundation grants from Stanley (A.S.), RUSK/S-R (A.S.), and NARSAD (A.S. and S.K.).

## References

1. Frith CD, Frith U: Mechanisms of Social Cognition. Annu Rev Psychol 2012; 63:287–313

2. Rilling JK, Sanfey AG: The Neuroscience of Social Decision-Making. Annu Rev Psychol 2011; 62:23–48

3. Bora E, Pantelis C: Social cognition in schizophrenia in comparison to bipolar disorder: A meta-analysis [Internet]. Schizophr Res 2016; 175:72–78 Available from: http://dx.doi.org/10.1016/j.schres.2016.04.018

4. Fett AKJ, Shergill SS, Krabbendam L: Social neuroscience in psychiatry: Unravelling the neural mechanisms of social dysfunction. Psychol Med 2015; 45:1145–1165

5. McCleery A, Lee J, Joshi A, et al.: Meta-analysis of face processing event-related potentials in schizophrenia [Internet]. Biol Psychiatry 2015; 77:116–126 Available from: http://dx.doi.org/10.1016/j.biopsych.2014.04.015

6. Meyer-Lindenberg A, Tost H: Neural mechanisms of social risk for psychiatric disorders. Nat Neurosci 2012; 15:663–668

7. Weightman MJ, Air TM, Baune BT: A review of the role of social cognition in major depressive disorder. Front Psychiatry 2014; 5:179

8. Gilboa-Schechtman E, Shachar-Lavie I: More than a face: a unified theoretical perspective on nonverbal social cue processing in social anxiety. Front Hum Neurosci 2013; 7:904

9. Kohler CG, Walker JB, Martin EA, et al.: Facial emotion perception in schizophrenia: A meta-analytic review. Schizophr Bull 2010; 36:1009–1019

10. Bicks LK, Koike H, Akbarian S, et al.: Prefrontal cortex and social cognition in mouse and man. Front Psychol 2015; 6:1805

11. Wilson CA, Koenig JI: Social interaction and social withdrawal in rodents as readouts for investigating the negative symptoms of schizophrenia [Internet]. Eur Neuropsychopharmacol 2014; 24:759–773 Available from: http://dx.doi.org/10.1016/j.euroneuro.2013.11.008

12. Craig AD: Once an island, now the focus of attention. Brain Struct Funct 2010; 214:395–396

13. Craig AD: How do you feel - now? The anterior insula and human awareness. Nat Rev Neurosci 2009; 10:59–70

14. Gogolla N: The insular cortex. Curr Biol 2017; 27:R580–R586

15. Phelps EA, Lempert KM, Sokol-Hessner P: Emotion and decision making: Multiple modulatory neural circuits. Annu Rev Neurosci 2014; 37:263–287

16. Downar J, Blumberger DM, Daskalakis ZJ: The Neural Crossroads of Psychiatric Illness: An Emerging Target for Brain Stimulation. Trends Cogn Sci 2016; 20:107–120

17. Singer T, Critchley HD, Preuschoff K: A common role of insula in feelings, empathy and uncertainty. Trends Cogn Sci 2009; 13:334–340

18. Namkung H, Kim SH, Sawa A: The Insula: An Underestimated Brain Area in Clinical Neuroscience, Psychiatry, and Neurology [Internet]. Trends Neurosci 2017; 40:200–207 Available from: http://dx.doi.org/10.1016/j.tins.2017.02.002

19. Uddin LQ: Salience processing and insular cortical function and dysfunction. Nat Rev Neurosci 2015; 16:55–61

20. Donaldson ZR, Young LJ: Oxytocin, vasopressin, and the neurogenetics of sociality. Science 2008; 322:900–904

21. Ferguson JN, Young LJ, Hearn EF, et al.: Social amnesia in mice lacking the oxytocin gene. Nat Genet 2000; 25:284–288

22. Rilling JK, Young LJ: The biology of mammalian parenting and its effect on offspring social development. Science (80-) 2014; 345:771–776

23. Knobloch HS, Charlet A, Hoffmann LC, et al.: Evoked axonal oxytocin release in the central amygdala attenuates fear response [Internet]. Neuron 2012; 73:553–566 Available from: http://dx.doi.org/10.1016/j.neuron.2011.11.030

24. Weissbourd B, Ren J, DeLoach KE, et al.: Presynaptic Partners of Dorsal Raphe Serotonergic and GABAergic Neurons [Internet]. Neuron 2014; 83:645–662 Available from: http://dx.doi.org/10.1016/j.neuron.2014.06.024

25. Jin D, Liu HX, Hirai H, et al.: CD38 is critical for social behaviour by regulating oxytocin secretion. Nature 2007; 446:41–45

26. Cataldo I, Azhari A, Esposito G: A review of oxytocin and arginine-vasopressin receptors and their modulation of autism spectrum disorder. Front Mol Neurosci 2018; 11:1–20

27. Febo M, Numan M, Ferris CF: Functional magnetic resonance imaging shows oxytocin activates brain regions associated with mother-pup bonding during suckling. J Neurosci 2005; 25:11637–11644

28. Bartels A, Zeki S: The neural correlates of maternal and romantic love. Neuroimage 2004; 21:1155–1166

29. Rogers CN, Ross AP, Sahu SP, et al.: Oxytocin- and arginine vasopressin-containing fibers in the cortex of humans, chimpanzees, and rhesus macaques. Am J Primatol 2018; 80:1–11

30. Dumais KM, Bredewold R, Mayer TE, et al.: Hormones and Behavior Sex differences in oxytocin receptor binding in forebrain regionsLJ: Correlations with social interest in brain region- and sex-speci fi c ways [Internet]. Horm Behav 2013; 64:693–701 Available from: http://dx.doi.org/10.1016/j.yhbeh.2013.08.012

31. Gehrlach DA, Dolensek N, Klein AS, et al.: Aversive state processing in the posterior insular cortex [Internet]. Nat Neurosci 2019; 22:1424–1437 Available from: http://dx.doi.org/10.1038/s41593-019-0469-1

32. Rogers-Carter MM, Varela JA, Gribbons KB, et al.: Insular cortex mediates approach and avoidance responses to social affective stimuli [Internet]. Nat Neurosci 2018; 21:404–414 Available from: http://dx.doi.org/10.1038/s41593-018-0071-y

33. Ng L, Bernard A, Lau C, et al.: An anatomic gene expression atlas of the adult mouse brain. Nat Neurosci 2009; 12:356–362

34. Inoue, R., Mori H: Serine Racemase Knockout Mice: Neurotoxicity, Epilepsy, and Schizophrenia., in Yoshimura T., Nishikawa T. HH, editor D-Amino Acids. 2016, pp 119–136.

35. Steinert JR, Chernova T, Forsythe ID: Nitric Oxide Signaling in Brain Function, Dysfunction, and Dementia. Neuroscientist 2010; 16:435–452

36. MacLean G, Abu-Abed S, Dollé P, et al.: Cloning of a novel retinoic-acid metabolizing cytochrome P450, Cyp26B1, and comparative expression analysis with Cyp26A1 during early murine development. Mech Dev 2001; 107:195–201

37. Catharine Ross A, Zolfaghari R: Cytochrome P450s in the Regulation of Cellular Retinoic Acid Metabolism. Annu Rev Nutr 2011; 31:65–87

38. Gandal MJ, Zhang P, Hadjimichael E, et al.: Transcriptome-wide isoform-level dysregulation in ASD, schizophrenia, and bipolar disorder. Science 2018; 362:eaat8127

39. Okano J, Lichti U, Mamiya S, et al.: Increased retinoic acid levels through ablation of Cyp26b1 determine the processes of embryonic skin barrier formation and peridermal development. J Cell Sci 2012; 125:1827–1836

40. Lalevée S, Anno YN, Chatagnon A, et al.: Genome-wide in Silico identification of new conserved and functional retinoic acid receptor response elements (direct repeats separated by 5 bp). J Biol Chem 2011; 286:33322–33334

41. Moutier E, Ye T, Choukrallah MA, et al.: Retinoic acid receptors recognize the mouse genome through binding elements with diverse spacing and topology. J Biol Chem 2012; 287:26328–26341

42. Saper CB: Convergence of autonomic and limbic connections in the insular cortex of the rat. J Comp Neurol 1982; 210:163–173

43. Uddin LQ, Nomi JS, Hébert-Seropian B, et al.: Structure and Function of the Human Insula. J Clin Neurophysiol 2017; 34:300–306

44. Swanson LW, Sawchenko PE: HYPOTHALAMIC INTEGRATIONLJ: Organization of the Paraventricular and Supraoptic Nuclei. Annu Rev Neurosci 1983; 6:269–324

45. Chang LJ, Smith A, Dufwenberg M, et al.: Triangulating the Neural, Psychological, and Economic Bases of Guilt Aversion [Internet]. Neuron 2011; 70:560–572 Available from: http://dx.doi.org/10.1016/j.neuron.2011.02.056

46. Nomi JS, Schettini E, Broce I, et al.: Structural connections of functionally defined human insular subdivisions. Cereb Cortex 2018; 28:3445–3456

47. Yoshida M, Takayanagi Y, Inoue K, et al.: Evidence that oxytocin exerts anxiolytic effects via oxytocin receptor expressed in serotonergic neurons in mice. J Neurosci 2009; 29:2259–2271

48. Takayanagi Y, Yoshida M, Takashima A, et al.: Activation of Supraoptic Oxytocin Neurons by Secretin Facilitates Social Recognition [Internet]. Biol Psychiatry 2017; 81:243–251 Available from: http://dx.doi.org/10.1016/j.biopsych.2015.11.021

49. Oettl LL, Ravi N, Schneider M, et al.: Oxytocin Enhances Social Recognition by Modulating Cortical Control of Early Olfactory Processing [Internet]. Neuron 2016; 90:609–621 Available from: http://dx.doi.org/10.1016/j.neuron.2016.03.033

50. Tan Y, Singhal SM, Harden SW, et al.: Oxytocin receptors are expressed by glutamatergic prefrontal cortical neurons that selectively modulate social recognition. J Neurosci 2019; 39:3249–3263

51. Raam T, McAvoy KM, Besnard A, et al.: Hippocampal oxytocin receptors are necessary for discrimination of social stimuli [Internet]. Nat Commun 2017; 8:1–14 Available from: http://dx.doi.org/10.1038/s41467-017-02173-0

52. Yoshimura R, Kiyama H, Kimura T, et al.: Localization of oxytocin receptor messenger ribonucleic acid in the rat brain. Endocrinology 1993; 133:1239–1246

53. Kennett GA, Wood MD, Bright F, et al.: SE 242084, a selective and brain penetrant 5-HT(2C) receptor antagonist. Neuropharmacology 1997; 36:609–620

54. Séjourné J, Llaneza D, Kuti OJ, et al.: Social Behavioral Deficits Coincide with the Onset of Seizure Susceptibility in Mice Lacking Serotonin Receptor 2c. PLoS One 2015; 1–15

55. Li QH, Nakadate K, Tanaka-Nakadate S, et al.: Unique Expression Patterns of 5-HT2A and 5-HT2C Receptors in the Rat Brain during Postnatal Development: Western Blot and Immunohistochemical Analyses. J Comp Neurol 2004; 469:128–140

56. Santana N, Bortolozzi A, Serrats J: Expression of Serotonin 1A and Serotonin 2A Receptors in Pyramidal and GABAergic Neurons of the Rat Prefrontal Cortex. Cereb Cortex 2004; 1100–1109

57. Santana N, Artigas F: Expression of Serotonin 2C Receptors in Pyramidal and GABAergic Neurons of Rat Prefrontal Cortex: A Comparison with Striatum. Cereb Cortex 2017; 27:3125–3139

58. Paine TA, Swedlow N, Swetschinski L: Decreasing GABA function within the medial prefrontal cortex or basolateral amygdala decreases sociability [Internet]. Behav Brain Res 2017; 317:542–552 Available from: http://dx.doi.org/10.1016/j.bbr.2016.10.012

59. Yamamuro K, Bicks LK, Leventhal MB, et al.: A prefrontal–paraventricular thalamus circuit requires juvenile social experience to regulate adult sociability in mice [Internet]. Nat Neurosci 2020; 23:1240–1252 Available from: http://dx.doi.org/10.1038/s41593-020-0695-6

60. Austgen JR, Dantzler HA, Barger BK, et al.: 5-Hydroxytryptamine 2C receptors tonically augment synaptic currents in the nucleus tractus solitarii. J Neurophysiol 2012; 108:2292–2305

61. Qiu J, Xue C, Bosch MA, et al.: Serotonin 5-Hydroxytryptamine 2C Receptor Signaling in Hypothalamic Proopiomelanocortin NeuronsLJ: Role in Energy Homeostasis in Females. Mol Pharmacol 2007; 72:885–896

62. Hitti FL, Siegelbaum SA: The hippocampal CA2 region is essential for social memory. Nature 2014; 508:88–92

63. Okuyama T, Kitamura T, Roy DS, et al.: Ventral CA1 neurons store social memory. Science 2016; 353:1536–1541

64. Reay WR, Cairns MJ, Atkins JR, et al.: Polygenic disruption of retinoid signalling in schizophrenia and a severe cognitive deficit subtype. Mol Psychiatry 2020; 25:719–731

65. Reay WR, Cairns MJ: The role of the retinoids in schizophreniaLJ: genomic and clinical perspectives [Internet]. Mol Psychiatry 2020; 25:706–718 Available from: http://dx.doi.org/10.1038/s41380-019-0566-2

66. Maden M: Retinoic acid in the development, regeneration and maintenance of the nervous system. Nat Rev Neurosci 2007; 8

67. Aoto J, Nam CI, Poon MM, et al.: Synaptic Signaling by All-Trans Retinoic Acid in Homeostatic Synaptic Plasticity [Internet]. Neuron 2008; 60:308–320 Available from: http://dx.doi.org/10.1016/j.neuron.2008.08.012

68. Chen L, Lau AG, Sarti F: Synaptic retinoic acid signaling and homeostatic synaptic plasticity. Neuropharmacology 2014; 78

69. Yip PK, Wong L, Pattinson D, et al.: Lentiviral vector expressing retinoic acid receptor b 2 promotes recovery of function after corticospinal tract injury in the adult rat spinal cord. Hum Mol Genet 2006; 15:3107–3118

