## Supplemental information for "Anterior insula-associated social novelty recognition: orchestrated regulation by a local retinoic acid cascade and oxytocin signaling"

**Data Supplement for Kim, An et al.,**


- Supplementary Methods,
- Method References,
- Supplementary Figure Legends
- Supplementary Figures
- Supplementary Table

#### Supplementary Methods

**Animals.** Wild-type mice (C57BL/6J; stock # 000664), Tg(RARE-Hspa1b/lacZ)12Jrt/J mice (stock # 008477)(70) and Thy1-YFP-H Tg mice (stock # 003782)(71) were purchased from the Jackson Laboratory. *Cyp26B1*-floxed (*Cyp26B1$^{flox/flox}$*) mice(39) were obtained from Dr. Maria I. Morasso (NIAMS, NIH) and Dr. Hiroshi Hamada (RIKEN). Except when stated otherwise, male mice were used for all experiments. All mice were housed in a controlled environment (temperature; humidity; 12-h light/dark cycle) and given *ad libitum* access to food and water. All procedures were approved by the Institutional Animal Care and Use Committee at Johns Hopkins University.

**Viral plasmid preparation.** The pAAV-U6-CMV-ZsGreen plasmid was obtained from the UPenn Vector Core, and the ZsGreen was replaced with mCherry using NheI and NotI restriction enzyme sites. The mCherry fragment was prepared by polymerase chain reaction (PCR) (sense, 5'-CTG CTA GCT AGC GCC ACC ATG GTG AGC AAG GGC GAG GAG G; antisense, 5'-TCT AGA GTC GCG GCC GCT ACT TGT ACA GCT CGT CCA TGC). The resulting pAAV-U6-CMV-mCherry plasmid was used to generate pAAV-U6-shControl-CMV-mCherry and pAAV-U6-shCyp26B1-CMV-mCherry.

We synthesized two sets of oligonucleotides: shControl (sense, 5'-GAT CCC GAT CGA ATG TGT ACT TCG ATT CAA GAG ATC GAA GTA CAC ATT CGA TCT TTT TTA AGC TTG; antisense, 5'-AAT TCA AGC TTA AAA AAG ATC GAA TGT GTA CTT CGA TCT CTT GAA TCG AAG TAC ACA TTC GAT CGG) and shCyp26B1 (sense, 5'-GAT CCG ATC CAA CTG GTG ATC CAG TTC AAG AGA CTG GAT CAC CAG TTG GAT CTT TTT TAA GCT TG; antisense, 5'-AAT TCA AGC TTA AAA AAG ATC CAA CTG GTG ATC CAG

TCT CTT GAA CTG GAT CAC CAG TTG GAT CG). The oligonucleotides had BamHI and EcoRI overhangs to allow for ligation into the pAAV-U6-CMV-mCherry. All final clones were verified by sequencing.

The mouse Cyp26B1 full-length cDNA plasmid (pYX-ASC-Cyp26B1) was obtained from OpenBiosystems. The human synapsin1 (hSyn) promoter-containing AAV plasmid (AAV-6P-SEWB) was a gift from Dr. Sebastian Kügler (Univ. of Göttingen); the EGFP of this plasmid was replaced by PCR-amplified mouse Cyp26B1-HA sequence to create pAAV-hSyn-Cyp26B1-HA plasmid using NheI and HindIII restriction enzyme sites: Cyp26B1-HA (sense, 5'-GCA CAG GCT AGC ACC GCG GTG GCG GCC GCC ATG CTG TTT GAA GGC TTG GAG; antisense, 5'-CCC CAC CCA AGC TTC TAT GCG TAG TCA GGC ACA TCA TAG GGG TAC ACC GTA GCA CTC AAC ATG GCC TCT GTC).

To create a RNAi-resistant Cyp26B1-expressing AAV vector, RNAi-target sequences of Cyp26B1 (5'-GAT CCA ACT GGT GAT CCA G) in the pAAV-hSyn-Cyp26B1-HA plasmid were changed using a site-directed mutagenesis kit (Agilent): sense, 5'-CTT GAG AGC TAC CTG CCC AAG ATT CAG TTA GTG ATC CAG GAT ACA CTT CG; antisense, 5'-CGA AGT GTA TCCT GGA TCA CTA ACT GAA TCT TGG GCA GGT AGC TCT CAA G; changed nucleotides are underlined and were verified by sequencing.

**Virus preparation.** The production of AAV was performed as described previously(72) with minor modifications. Human embryonic kidney (HEK) 293-AAV cells (Stratagene) were cotransfected with a transgene plasmid along with two packaging plasmids for AAV1 using a standard calcium phosphate precipitation method. Cells were harvested 72 h post transfection and disrupted by three freeze and thaw cycles. AAV viral particles were isolated and concentrated by a series of ammonium sulfate precipitations, applied to a discontinuous gradient of iodixanol (Optiprep™ density gradient medium, Sigma-Aldrich), and centrifuged at 350,000 x g for 60 min, 18 °C. An AAV-containing 40% iodixanol fraction was carefully collected from an ultracentrifuge tube (Beckman), and concentrated using a centrifugal filter device (Ultracel-100K, Amicon). Viral titers were finally adjusted to 1.2 x $10^{12}$–4.0 x $10^{12}$ particles/ml depending on experimental purposes. The AAV-αCaMKII-hChR2 (H134R)-EYFP, AAV-αCaMKII-GFP-Cre, and AAV-αCaMKII-EYFP were prepared in the viral vector core at the University of North Carolina.

**Cortical cell dissociation and fluorescence-activated cell sorting (FACS) purification.** Three groups of wild-type adult male mice (9 to 10-week-old) injected with corresponding AAV mixtures into their bilateral anterior insula (AI) were prepared for FACS: AAV-U6-shControl-CMV-mCherry and AAV-αCaMKII-EYFP; AAV-U6-shCyp26B1-CMV-mCherry and AAV-αCaMKII-EYFP; AAV-U6-shCyp26B1-CMV-mCherry, AAV-hSyn-Cyp26B1$^{R}$-HA and AAV-αCaMKII-EYFP. Mice were anesthetized with isoflurane, brains were removed and immediately sectioned in the coronal plane at 500 μm on a cold metal brain slicer (Zivic instrument), and slices were transferred into ice-cold Hank's Balanced Salt Solution (HBSS; Gibco). The AI was dissected from brain slices in ice-cold HBSS media and was further dissected to obtain highly enriched cortical layer 5 tissues. These dissected tissues were cut into small pieces and dissociated for 1 h at 37 °C with papain enzyme (Papain Dissociation System, Worthington Biochemical) in Earls Balanced Salt Solution (EBSS) with DNase according to the manufacturer's protocol. The tissues were gently triturated with three glass pipettes of decreasing tip diameter and dissociated cells were centrifuged at 300 x g for 5 min at room temperature. To remove excess cell debris, cell pellets were resuspended in EBSS containing DNase and albumin ovomucoid inhibitor (AOI), and the cell suspension was subjected to centrifugation on an AOI discontinuous gradient according to the manufacturer's protocol. Cell pellets were resuspended in a HBSS media containing 1% bovine serum albumin (BSA), filtered through a 40 µm mesh (BD Falcon), and sorted on a FACS Aria II cell sorter (Becton Dickinson) by the expression of EYFP and mCherry. Double-positive cells were directly sorted into a 1.5 ml tube containing RNA extraction buffer.

**Gene expression profiling.** RNAs were extracted using the PicoPure RNA Isolation kit (Arcturus) with a DNase I digestion according to the manufacturer's protocol. RNA quality and relative concentration were examined on the Picochip using an Agilent 2100 Bioanalyzer (Agilent Technologies). The RNA integrity number (RIN) values of all the RNA samples used in further analyses were greater than 7.4, with an average RIN value of 8.3. The RNA samples (500 pg per sample) were amplified using the Pico WTA V2 system (Nugen) and the amplified cDNA was labelled using Encore Biotin Module Nugen (Nugen) following the manufacturer's recommended procedure. The labelled product was then hybridized to an Affymetrix mouse Gene 2.0ST array at 45 °C with 60 rpm rotation for 18 h, then washed and stained using an Affymetrix wash and stain kit and scanned using a GeneChip G3000 scanner (Affymetrix). Hybridization, washing and scanning were conducted according to the manufacturer's instructions. All microarray procedures were carried out at the Microarray Core Facility of Johns Hopkins University.

**Quantitative real-time PCR (qRT-PCR).** cDNA synthesis was performed using the SuperScript™ First-Strand System for RT-PCR (Invitrogen) and Oligo(dT)$_{20}$ primers according to the manufacturer's instructions. Specific primers for the target mouse genes were designed by Primer-blast (NCBI). Primer sequences are as follows: Sytl2 (sense, 5'-ACA CAG CAT CAG AGA GCA GC; antisense, 5'-ACA AGG ACG TCA TGC CAG AG); Ephb2 (sense, 5'-ATG TAA CAG ACG GGG GTT TGA G; antisense, 5'-GGT CAT GTG TCC GCT GGT GT); Dip2a (sense, 5'-TCT GCC TCC AGG GCA CAA CA; antisense, 5'-ATG CGG AGA ACC CCG TTC CA); β-actin (sense, 5'-AGG CTC TTT TCC AGC CTT CC; antisense, 5'- CAT GGA TGC CAC AGG ATT CC); Gapdh (sense, 5'-CGT GGG GCT GCC CAG AAC AT; antisense, 5'-CAG GCG GCA CGT CAG ATC CA). qRT-PCR was carried out using SYBR® GreenER™ qPCR SuperMix for ABI PRISM® (Invitrogen) for the 7900HT Sequence Detector System (Applied Biosystems) in a final volume of 10 μl. The PCR conditions for 7900HT were set as: 50°C for 2 min, 95 °C for 10 min followed by 40 cycles of 95 °C for 15 s, 60 °C for 1 min and a dissociation step of 95 °C for 15 s, 60°C for 15 s, and 95 °C for 15 s. The expression level of β-Actin and GAPDH genes were used for normalization. We analyzed the data with SDS 2.4 (Applied Biosystems).

**Stereotaxic injections.** Adult mice (9 to 10-week-old) were deeply anesthetized with a mixture of ketamine/xylazine (ketamine, 100 mg/kg; xylazine, 10 mg/kg; intraperitoneal injection) and placed into a stereotaxic apparatus (David Kopf Instruments). Neurotoxic lesions were made by injecting 0.15 μl of *N*-methyl-D-aspartate [NMDA; 15 mg/ml in phosphate buffered saline (PBS), Sigma] at two sites per hemisphere using a glass micropipette controlled by an auto-nanoliter injector (Nanoject II, Drummond). The following coordinates from bregma were used -

orbitofrontal cortex (OFC): +2.4 AP, ±1.2 ML, −1.75 DV and +2.0 AP, ±1.6 ML, −2.3 DV; AI: +2.4 AP, ±2.15 ML, −1.8 DV and +2.0 AP, ±2.65 ML, −2.0 DV. Using the same coordinates, 0.3 µl (per hemisphere) of viral solutions or 0.2 µl of retrograde tracer, cholera toxin subunit B-conjugated with Alexa 488 (CTB-Alexa 488, 5 µg/µl in PBS, Invitrogen; right hemisphere) was injected. After each injection of drugs, virus, or tracer, the micropipette was left in place for 10 min and then slowly withdrawn. The skin was sutured closed. Mice fully recovered under a heat lamp before returning to their home cage and were used for corresponding behavioral tests 4 weeks later.

**Cannula or optic fiber implantation.** Twenty-six gauge guide cannulae (Plastics One) were implanted bilaterally over the AI (+2.0 AP, ±2.60 ML, −0.5 DV), unilaterally into the right lateral cerebral ventricle (−0.3 AP, +0.8 ML, −0.5 DV), or dorsal raphe nucleus (DRN) (−4.2 AP, 0 ML, −3.0 DV) at a 30° angle. The cannulae were secured in place with anchoring miniature screws and dental cement. A 32-gauge dummy cannula (Plastics One) was inserted into each cannula to prevent clogging. The cannula positions were verified in DiI (Invitrogen)-stained cryosections.

Mice used in optogenetic stimulation experiments received virus injection and optic fiber surgery on the same day. After virus injection, optic fibers [200 nm core diameter, 0.22 numerical aperture (NA), Thorlabs] were implanted bilaterally over the injection site (+2.2 AP, ±2.40 ML, −1.5 DV). The light out through the bilateral optic fiber was measured prior to bilateral implantation. Fibers measuring at least 10 mW were utilized. Optic fibers were secured to the skull with dental cement (Lang dental) and two miniature screws attached them to the skull.

***In vivo* optical stimulation.** During behavioral testing, an external optical fiber (62.5 nm core diameter, 0.22 NA, Precision Fiber Products) was coupled with the implanted fiber optic using a zirconia sleeve. An optical commutator (Doric Lenses) allowed for unrestricted rotation. Optical stimulation was provided with a 100 mW 473 nm diode pumped solid state laser (Laserglow Technologies) and controlled by a Master 8 pulse generator (A.M.P.I.). During the trial 1 (10 min) and trial 2 (10 min) sessions in the three-chamber social interaction test, blue laser optical stimulation (473-nm, 5-Hz, 10-ms duration; around 10 mW output at the fiber tip) was consistently delivered to mice.

**Drug treatment.** For drug delivery through cannulas, the dummy cannulas were removed and injections were made through a 33-gauge internal cannula (Plastics One) extending 2 mm below the tip of the guide cannula. The internal cannula was backfilled with mineral oil (Sigma) and connected to a 25-μl syringe (Hamilton). Depending on the experimental purpose, the following drugs were delivered into the AI, intracerebroventricular (i.c.v.), or DRN, respectively. For chronic retinoic acid (RA) infusion, either RA [2 mg/ml dissolved in 50% dimethyl sulfoxide (DMSO)] or vehicle (50% DMSO) was injected at a rate of 100 nl/min for 1 min per day for 16 consecutive days using a microinfusion pump (Narishige). The internal cannula was left in place for an additional 5 min to achieve a proper diffusion of the drug or vehicle. Eighteen h after the completion of chronic drug injections, mice performed the three-chamber social interaction test. For acute local administration, 5-HT2C receptor agonist (CP 809101, 3.4 ng/100 nl, Tocris) or its antagonist (SB 242084, 50 ng/μl, Tocris), OT (2.5 μg/μl, Tocris) or OTA (desGly-$NH_2$-d$(CH_2)_5$ [D-$Tyr^2$, $Thr^4$] OVT, 50 ng/μl, Bachem) dissolved in artificial cerebrospinal fluid (ACSF) were infused into their corresponding brain regions at a rate of 0.1 µl/min for 1 min and waited for an additional 10 min. For i.c.v. infusion, OT (1 ng/µl) or OTA (20 ng/µl) was infused at a rate of 0.25 µl/min for 20 min and waited for an additional 10 min. For the systemic treatment, 5-HT1A receptor agonist (Tandospirone, 1 mg/kg, Tocris) or its antagonist (WAY 100635, 0.6 mg/kg, Tocris), and CP 809101 (1 mg/kg) or SB 242084 (1 mg/kg) dissolved in PBS were subcutaneously (sc) injected. Mice performed the behavioral test 30 min after the completion of the drug treatment.

***In situ* hybridization (ISH).** The Cyp26B1 sequences for the ISH probe were prepared by PCR (sense, 5'-CGC TCA GCA GCG GCC GCT ACC TGG ACT GTG TCA TCA AG; antisense, 5'-CCT CCC CTC GAG ACA GGG ATC CCC TTC AGC TTT TCC; start from 1057 nt of ORF), and were inserted into NotI and XhoI sites of the pBluescript SK (Stratagene) plasmid. Plasmid DNA for probe synthesis was linearized with restriction enzyme, checked for completeness of digestion on agarose gel, and then protease K-digested, phenol/chloroform-extracted three times, and collected via ethanol precipitation. Probes were synthesized with T3 or T7 polymerase and checked on agarose gel. Sixteen µm-thick mouse brain cryosections were hybridized with digoxigenin-labeled probes (Roche) at 70 °C overnight. The next day, excess probes were washed out and sections were blocked with 10% sheep serum and incubated with buffer solution containing alkaline phosphatase-conjugated antibodies (Roche) at 4 °C overnight. Color was developed with the developing solution [100 mM Tris pH9.8, 100 mM NaCl, 50 mM $MgCl_2$, 5% polyvinyl alcohol, 0.11 mM 5-bromo-4-chloro-3'-indolyphosphate (BCIP), 0.12 mM nitroblue tetrazolium (NBT), and 0.5 mg/ml levamisol; all purchased from Sigma-Aldrich] at 37 °C. The reaction was stopped in water. Sections were dehydrated through an ascending alcohol series

(30, 50, 70, 90, and 100% ethanol) followed by xylene, and mounted with Permount (Fisher) mounting medium.

**X-gal staining.** The reporter mice for RA signaling [Tg(RARE-Hspa1b/lacZ)12Jrt/J] injected with corresponding AAVs into the AI were briefly perfused transcardially with 2% glutaraldehyde, and the removed brains were embedded in low-melting point agarose and sliced at 200 μm on a vibratome (VT1200S, Leica). The slices were collected in 24-well plates and processed for lacZ visualization in X-gal-containing color reaction buffer at 37 °C. The reaction was terminated in 4% paraformaldehyde at 4 °C, rinsed in PBS, and mounted with aqueous mounting medium (DAKO).

**Quantitative dendritic spine analysis.** Three groups of adult (9 to 10-week-old) Thy1-YFP-H Tg mice injected with corresponding AAVs into their bilateral AI were prepared for dendritic spine analysis: Control, AAV-U6-shControl-CMV-mCherry; knockdown (KD), AAV-U6-shCyp26B1-CMV-mCherry; Rescue, AAV-U6-shCyp26B1-CMV-mCherry and AAV-hSyn-Cyp26B1$^{R}$-HA. After perfusion and fixation with 4% paraformaldehyde, coronal sections (50 μm thickness) were obtained by the vibratom. The slices were further stained with anti-GFP antibody (1:500, Abcam) and AlexaFluor 488-conjugated secondary antibody (1:400, Life Technology) to circumvent potential unevenness of YFP diffusion in spines. Images were taken by a Zeiss LSM510-Meta with the 63x oil-immersion objective as *z* series of 6–12 images taken at 0.3 μm intervals (scan averaged, four times; 1024 x 1024 pixel resolution; a scan speed of 1.6 ms per pixel). The acquisition parameters were kept constant for all scans in the same experiment. Intact YFP-positive pyramidal neurons within layer 5 of the AI were selected for imaging based on the following criteria: (i) Location of cell soma within layer 5 of the AI (AP +2.34 to +1.98), (ii) YFP and mCherry double-positive cells, (iii) Healthy neurons with no signs of blebbing/distress, (iv) Intact dendritic arbor with primary apical dendrite that extends to layer 2/3 without truncation. For these neurons, secondary and tertiary dendrites that branched from an apical dendrite between 50 μm and 200 μm from the cell soma were selected for spine analysis. Branch segments of 30 μm without additional branching were identified and analyzed for spine density. Individual spines (>3 μm in length) on dendrites were manually traced and analyzed using MetaMorph software version 7.1 (MDS Analytical Technologies). Spine density was quantified as the number of spines per 10 μm dendritic segment. We analyzed more than 500 spines derived from 13 to 15 neurons per condition. The analyses were carried out while blinded to the identity of each condition until statistical analysis had been done.

**Electrophysiology.** Acute brain slices were dissected as previously described(73). Briefly, mice were deeply anesthetized with isoflurane and then transcardially perfused with ice-cold oxygenated (95% $O_2$ and 5% $CO_2$) dissecting buffer containing (in mM): 83 NaCl, 2.5 KCl, 1 $NaH_2PO_4$, 26.2 $NaHCO_3$, 22 dextrose, 72 sucrose, 0.5 $CaCl_2$, and 3.3 $MgCl_2$, adjusted to pH 7.3. The brain was rapidly removed and immersed in ice-cold oxygenated dissection buffer. Coronal slices (300 μm thickness) were cut using the vibratome, incubated in dissection buffer for 30–45 min at 34 °C, and then kept at room temperature in ACSF containing (in mM): 125 NaCl, 25 $NaHCO_3$, 1.25 $NaH_2PO_4$, 3 KCl, 25 dextrose, 1 $MgCl_2$, and 2 $CaCl_2$, adjusted to pH 7.3. Slices were visualized by IR DIC microscopy (BX51WI, Olympus) and a CCD camera (QICAM, QImaging). Individual neurons expressing fluorescent proteins were visualized with epifluorescent illumination and a 40 x Olympus LUMPlan FLN water immersion (0.8 NA) objective. The recording electrodes (3–5 MΩ) were filled with internal solution containing (in mM): 125 potassium gluconate, 10 HEPES, 4 Mg-ATP, 0.3 Na-GTP, 0.1 EGTA, 10 phosphocreatine, adjusted to pH 7.3 with KOH. Whole-cell patch-clamp recordings were obtained with a Multiclamp 700B amplifier (Molecular Devices) controlled by pClampex10.3 (Axon Instruments). The output signals were filtered at 2 kHz using built-in Bessel filters and digitized at a sampling rate of 20 kHz with Digidata 1440A (Molecular Devices). Spontaneous excitatory postsynaptic currents (sEPSCs) were obtained with SR95531 hydrobromide (gabazine, 5 μM, Tocris) in ACSF while clamping membrane potential at -70 mV. Only results from the cells with stable access resistance (change less than 20%) were used for further analysis. The data were displayed off-line with Clampfit software (Axon Instruments), EPSCs were detected automatically with MINIANALYSIS (Synaptosoft) and each event was manually inspected. To compare paired-pulse ratios (PPR), evoked EPSCs were triggered with a bipolar concentric stimulating electrode (FHC) placed at the superficial layer of the AI. Stimulating intensity was adjusted to induce 100–300 pA of EPSCs at -70 mV. PPRs were acquired by dividing EPSC2 with EPSC1 obtained by consecutive stimulation with 50, 100, 200, and 400-ms inter-stimulus intervals. For pharmacological examinations, the recordings were conducted following more than 10 min of 50 μM of CP 809101 or 10 μM of SB 242084 exposure.

**Behavioral assays.** All tests were conducted following the protocols as described during the light cycle. In between animals, all testing chambers and apparatuses were cleaned with 70% ethanol and fully dried. Unless otherwise stated, mice behaviors were monitored with an Ethovison XT (Noldus) tracking system and manually analyzed in a blinded manner.

***Three-chamber social interaction.*** The three-chamber social interaction test(74) arena consisted of three adjacent chambers separated by two clear plastic dividers and connected by two open doorways. Mice were habituated for 3 consecutive days (10 min per day) before a testing day. Mice were transferred to a testing room and stayed for 1 h before testing. The test consisted of 5 min of habituation and two 10 min sessions (trial 1 and 2) with a 5-min inter-trial interval. The subject mouse was kept in a center chamber during habituation. In the first 10 min session (trial 1), the subject mouse freely investigated the three chambers and a novel object mouse (stranger 1) was placed under an inverted metal cup in one of the side chambers and another identical inverted empty cup was placed in the other side chamber. After the trial 1 session, the doorways were closed and the subject mouse was kept in the center chamber for 5 min. In the second session (trial 2), another novel object mouse (stranger 2) was put in the empty cup and the subject mouse freely investigated the three chambers. Age-matched adult C57BL/6J male mice (Jackson Laboratory) were used as object mice in this study, and housed under the same conditions as the subject mice. Each object mouse was used once a day in the three-chamber social interaction test. Time spent sniffing each cup was scored as interaction time.

***Five-trial recognition test.*** For the 5-trial recognition test with social cues, we transferred mice from group to individual housing for 7–10 days before testing to permit the establishment of a home-cage territory. The test began when a stimulus female mouse was introduced into the home cage of each male mouse for a 1 min confrontation. Age-matched ovariectomized female C57BL/6 mice (Taconic) were used as stimulus mice. At the end of the 1 min trial, the stimulus mouse was removed and returned to an individual holding cage. This sequence was repeated for four trials with 10-min inter-trial intervals. The same stimulus mouse was introduced to the same male resident in all four trials. In a fifth dishabituation trial, we introduced a different stimulus mouse to the resident male mouse. Behavior was recorded and interaction time (time for nosing, anogenital sniffing, and close following and pursuit) was scored. Aggressive posturing and sexual behaviors (e.g. mounting) were not included in the calculation of social interaction time. For the 5-trial recognition test with non-social cues, we transferred mice from group to individual housing for 7–10 days before testing. A cotton ball with one microliter of natural banana oil (Lorann oils) was placed in a perforated 15 ml conical tube with eight 2 mm-diameter holes near the ball, and the scented tube was placed in the home cage of each mouse for 1 min. We repeated this four times with 10 min intervals. In a fifth trial, we placed a 15 ml conical tube containing a cotton ball with the same volume of natural strawberry oil (Lorann oils) in each subject cage for 1 min. We operationally defined olfactory investigation as nasal contact with the tube.

***Olfactory and locomotor test.*** We used the hidden food test to examine the basic olfactory function of mice: the mice were food-deprived for 24 h and were habituated to a new cage for 5 min before testing. After cheese (0.5 cm x 0.5 cm x 0.5 cm) was buried beneath 2 cm of leveled bedding, each subject mouse was put into the cage and the latency to locate the cheese was recorded. We used the open field test to examine the basic locomotor activity of mice: each mouse was placed in a novel open field box (40 cm x 40 cm; San Diego Instruments) for 1 h. Horizontal and vertical locomotor activities were automatically recorded by an infrared activity monitor (San Diego Instruments). Single beam breaks are reported as “counts”.

**Bioinformatics.**

***Genome-wide association studies (GWAS) analysis.*** A list of genes that are enriched in the AI layer 5 pyramidal neurons were selected by comparing a publicly available database from the

Allen Institute for Brain Science [anatomic gene expression analysis (AGEA)](33) with our microarray database acquired from the gene expression profiling. Genes enriched in the AI layer 5 were defined as a fold change above 1 by comparing expression levels between the region of interest and other brain regions (AGEA database), and were inspected for their expression in the pyramidal neurons (microarray intensity > 5). The resulting 1205 genes were subjected to further analysis by GWAS for schizophrenia using data from NHGRI-EBI GWAS Catalog(75) to identify disease-relevant risk genes (odds ratio ≥ 1 and –log (*P*-value) ≥ 6). The list of genes identified as risk genes from this analysis were categorized into 7 groups based on their specific functions (Extended Data Fig. 3a).

***Signal pathway analysis for brain disorders.*** An enrichment analysis on 215 signal transduction pathways from GO biological process was performed to identify over-represented pathways in risk genes for brain disorders. The risk genes for brain disorders are genes with 1 or more single nucleotide polymorphisms (SNP) associated with brain disorders [schizophrenia, bipolar disorder, psychosis, Alzheimer's disease, Parkinson's disease, dementia, attention-deficit/hyperactivity disorder (ADHD), autism, cognitive decline, and depression] (*P*-value < 1E-6) from the NHGRI-EBI GWAS Catalog(75). Over-representation was assessed using a chi-square test for the 2 x 2 contingency table (Inside-/outside-targeted pathway x risk/not-risk gene).

***Molecular Profiling data analysis.*** Data analysis was performed using the Partek Genomics Suite software (version 6.6, Partek Inc., St Louis, MO, USA). Partek default settings were used for data import and normalization; raw intensities were normalized using robust multi-array average background correction with adjustment for GC-content. Principal components analysis was performed using a correlation matrix. An ANOVA model examining the treatment factor with contrasts for Cyp26B1 KD vs. Control, Cyp26B1 KD vs. Rescue, and Control vs. Rescue, was used to determine transcripts that were significantly regulated. The ANOVA *P*-values were adjusted using the Benjamini–Hochberg procedure to control the false discovery rate (FDR). Transcripts with no RefSeq identifier and/or no gene symbol were removed. In order to gain a broad overview of the effect of knocking down Cyp26B1, the 114 transcripts (corresponding to 108 unique genes) that survived FDR 0.2 with a minimum fold-change of |1.2| were used to assess the presence of RAREs and enrichment of gene ontology (GO) terms. Partek GO-Enrichment analysis was conducted using the population of genes present in the Affymetrix annotation file. The 108 unique genes were examined for overlap with the mouse and human RARE lists from Lalevée and coworkers(40). Hierarchical clustering (Partek) was conducted on the 20 genes that survived FDR 0.1 with a minimum fold-change of |1.4|.

#### Method References


70. Rossant J, Zirngibl R, Cado D, et al.: Expression of a retinoic acid response element-hsplacZ transgene defines specific domains of transcriptional activity during mouse embryogenesis. Genes Dev 1991; 5:1333–1344

71. Feng G, Mellor RH, Bernstein M, et al.: Imaging Neuronal Subsets in Transgenic Mice Expressing Multiple Spectral Variants of GFP [Internet]. Neuron 2000; 28:41–51Available from: http://linkinghub.elsevier.com/retrieve/pii/S0896627300000842

72. Choi VW, Asokan A, Haberman RA, et al.: Production of Recombinant Adeno-Associated Viral Vectors for In Vitro and In Vivo Use. Curr Protoc Mol Biol 2007; 78:16.25.1-16.25.24

73. Maher BJ, LoTurco JJ: Disrupted-in-schizophrenia (DISC1) functions presynaptically at glutamatergic synapses. PLoS One 2012; 7:1–9

74. Nadler JJ, Moy SS, Dold G, et al.: Automated apparatus for quantitation of social approach behaviors in mice. Genes, Brain Behav 2004; 3:303–314

75. Buniello A, Macarthur JAL, Cerezo M, et al.: The NHGRI-EBI GWAS Catalog of published genome-wide association studies, targeted arrays and summary statistics 2019. Nucleic Acids Res 2019; 47:D1005–D1012

#### Supplement Figure Legends

**FIGURE S1. Excitotoxic lesions in the OFC and AI.**

**A,** Coronal schematics of the AI and PI to indicate that the AI regions are distinct from the PI. Blue and red indicate the AI and PI, respectively.

**B**, Coronal schematics of NMDA-induced brain lesions. The representative lesions are indicated in blue. OFC, n = 3; and AI, n = 7.

**FIGURE S2. Behavior in OFC- and AI-lesioned mice.**

**A**, Normal olfactory function in OFC- (left) and AI-lesioned (right) mice. OFC-Con, n = 5; OFC-lesion, n = 5; AI-Con, n = 7; and AI-lesion, n = 7.

**B,** Normal locomotor activity in OFC- (left) and AI-lesioned (right) mice. OFC-Con, n = 5; OFC-lesion, n = 5; AI-Con, n = 10; and AI-lesion, n = 9.

**C**, No gender difference in the social interaction behavioral paradigm in OFC- (left) and AI-lesioned (right) mice. Female mice exhibit the same pattern as male mice in the behavioral paradigm. OFC-Con, n = 5; OFC-lesion, n = 8; AI-Con, n = 10; and AI-lesion, n = 10.

Two-tailed *t*-test for **A** and **B**. Two-way ANOVA with Bonferroni *post-hoc* test for **c**. Data are represented as mean ± S.D.

*** $p<0.001$.

**FIGURE S3. Bioinformatic analysis for the selection of Cyp26B1.**

**A,** Forty seven candidate molecules selected with a bioinformatic approach: Cyp26B1 was one of two molecules in the “enzymes critical to biosynthesis of key mediators” class. Class 1, receptor/transporter; class 2, ligand/cell adhesion molecule; class 3, transcriptional and nuclear factor; class 4, signal transduction; class 5, cytoskeleton-related; class 6, enzymes critical to biosynthesis of key mediators; and class 7, unclassified.

**B**, RA-signaling related genes highlighted in a transcriptome analysis for autism spectrum disorder, schizophrenia, and bipolar disorder. Genes highlighted in red indicate a significant increase in expression, while genes highlighted in blue indicate a significant decrease in expression. Only Cyp26B1 and PTK2B were significant in all three conditions: Cyp26B1 was consistently decreased in all conditions, whereas PTK2B was upregulated in schizophrenia and bipolar disorder (↑) and downregulated in autism spectrum disorder (↓).

**FIGURE S4. Histological validation of the surgical manipulation of RA delivery or AAV injection.**

**A-C**, Coronal schematics of the affected mouse brain areas at the indicated distance from the bregma. Red and blue circles (**A**): location of injection cannula tips. Blue: representative spread of fluorescent dye DiI (**A**), mCherry (**B**) and GFP (**C**) expressed by infected AAV.

**FIGURE S5. The behavioral and molecular effect of Cyp26B1 KD in the AI.**

**A**, Representative images of i*n situ* hybridization for Cyp26B1 (violet) in control (Con) and AI-Cyp26B KD (KD). Scale bar, 500 μm.

**B**, Representative images of β-galactosidase color reaction (green) with the RARE-LacZ reporter (indicator of RA signaling) in control (Con) and AI-Cyp26B KD (KD). Scale bar, 1 mm.

**C**, AI-Cyp26B1 KD mice exhibit selective deficits in social recognition but exhibit normal performance in the 5-trial recognition test with non-social cues compared with controls (Con). Con, n = 8; and KD, n = 8.

**D**, qRT-PCR validation of RA-regulated genes. AAV-Con (Con); KD (Cyp26B1 KD); and KD + Cyp26B1$^R$ (Rescue). n = 3 per group.

Two-way repeated measure ANOVA with Bonferroni *post-hoc* test for **C**; one-way ANOVA with *post-hoc* Tukey's multiple comparison for **D**. Data are represented as mean ± S.D. *$p<0.05$, ** $p<0.005$, *** $p<0.001$.

**FIGURE S6. Histological and electrophysiological validation of the optogenetic manipulation of AI neurons.**

**A**, Coronal schematics indicating the placement of optic fibers and AAV expression. Blue and red circles: location of optic fiber tips. The representative extent of EYFP expression is indicated in blue.

**B**, Whole cell recording from an AI layer 5 pyramidal neuron co-expressing mCherry and ChR2-EYFP in a slice from a mouse injected with Cyp26B1 KD AAV (mCherry) and ChR2-EYFP AAV. Activation with a 473-nm laser pulse (blue light) for 10-ms at 5-Hz (blue bars) triggered a non-adapting action potential in the recorded neurons. Scales, 1s and 10 mV. Inset shows expanded 1 s period with laser pulse stimulation. Inset scales, 100 ms and 20 mV.

**FIGURE S7. Histological validation of the surgical manipulations of drug infusion or AAV injection.**

**A, B**, Coronal schematics indicating the placement of cannula (blue circles) targeted to the AI (**A**) or i.c.v. (**B**). Blue: the extent of mCherry expression after injection of Control (Con) or Cyp26B1 KD (KD) AAV into the AI. Con AAV + Cannula (AI), n = 10; KD AAV + Cannula (AI), n = 12; Con AAV + Cannula (i.c.v.), n = 9; and KD AAV + Cannula (i.c.v.), n = 10.

**FIGURE S8. Pharmacological and electrophysiological studies to address 5-HT2C involvement in AI-mediated social novelty recognition.**

**A,** A pharmacological study to test the involvement of 5-HT1A and 5-HT2C receptors in AI-mediated social novelty recognition. Left: the 5-HT2C agonist CP 809101 (CP) (subcutaneously, s.c.) rescues the social novelty recognition deficits elicited by AI-Cyp26B1 KD ($n = 10$ per group). Right: the 5HT2C antagonist SB 242084 (SB) disrupts normal social recognition in control mice ($n = 9$ per group). Tan, Tandospirone (5-HT1A agonist); and Way, WAY 100635 (5-HT1A antagonist).

**B,** CP does not influence presynaptic vesicle release probabilities. A paired pulse ratio (PPR) was measured in Cyp26B1 KD (KD) AAV-infected neurons from AI brain slices. KD + Veh, $n = 9$ neurons from 4 mice; and KD + CP, $n = 9$ neurons from 4 mice.

Two-way ANOVA with Bonferroni *post-hoc* test. Data are represented as mean ± S.D.

*** $p<0.001$.

**FIGURE S9. Histological validation of the surgical manipulations of drug infusion into the DRN and AAV injection into the AI.**

**A, B**, Coronal schematics indicating the placement of cannula (blue circles) targeted to the DRN with the extent of mCherry expression after injection of Control (Con) or Cyp26B1 KD (KD) AAV into the AI. Con AAV + Cannula (DRN), $n = 10$; and KD AAV + Cannula (DRN), $n = 6$.

FIGURE S1

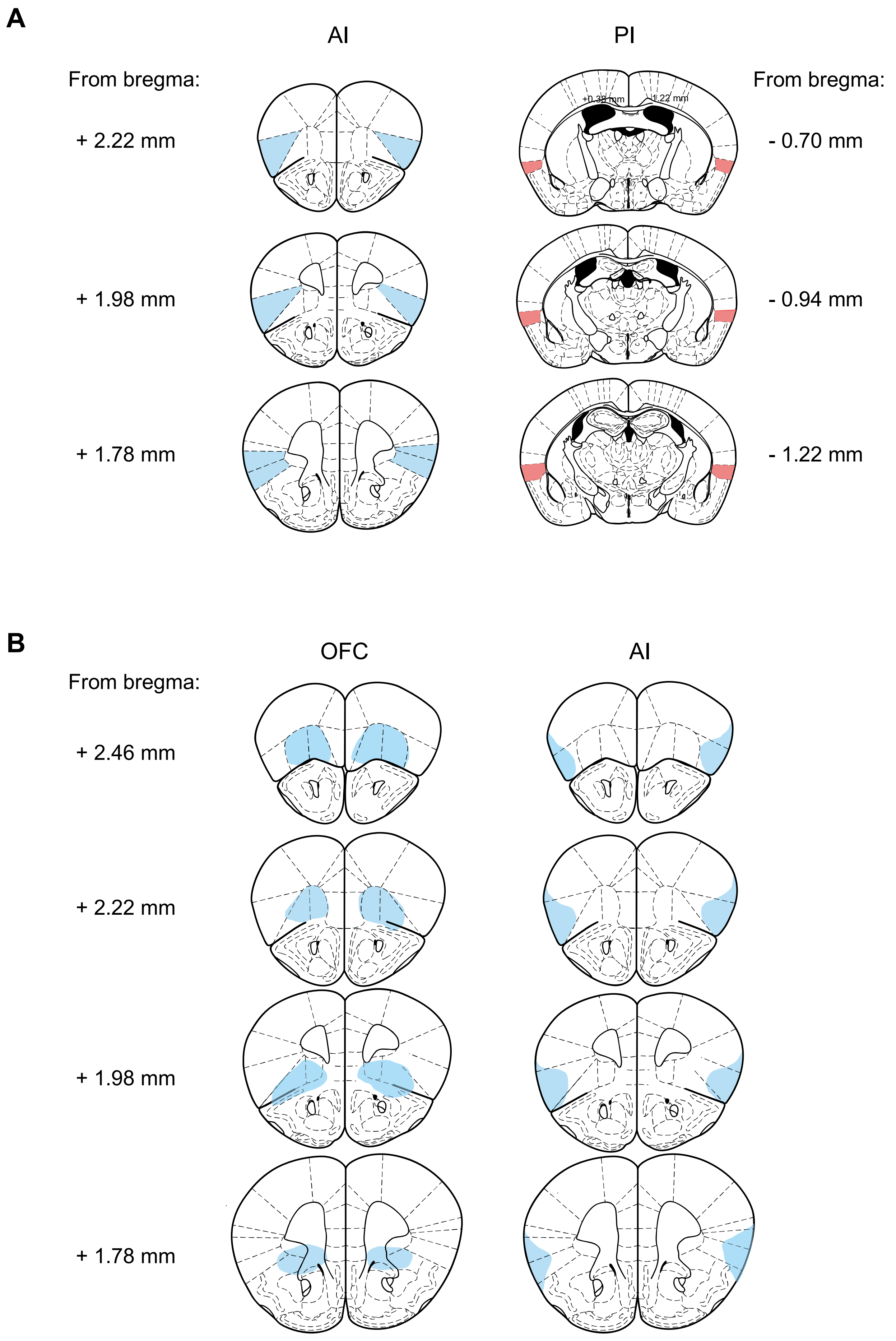

A
AI
PI
From bregma:
+ 2.22 mm
+ 1.98 mm
+ 1.78 mm
From bregma:
- 0.70 mm
- 0.94 mm
- 1.22 mm
B
OFC
AI
From bregma:
+ 2.46 mm
+ 2.22 mm
+ 1.98 mm
+ 1.78 mm

FIGURE S2

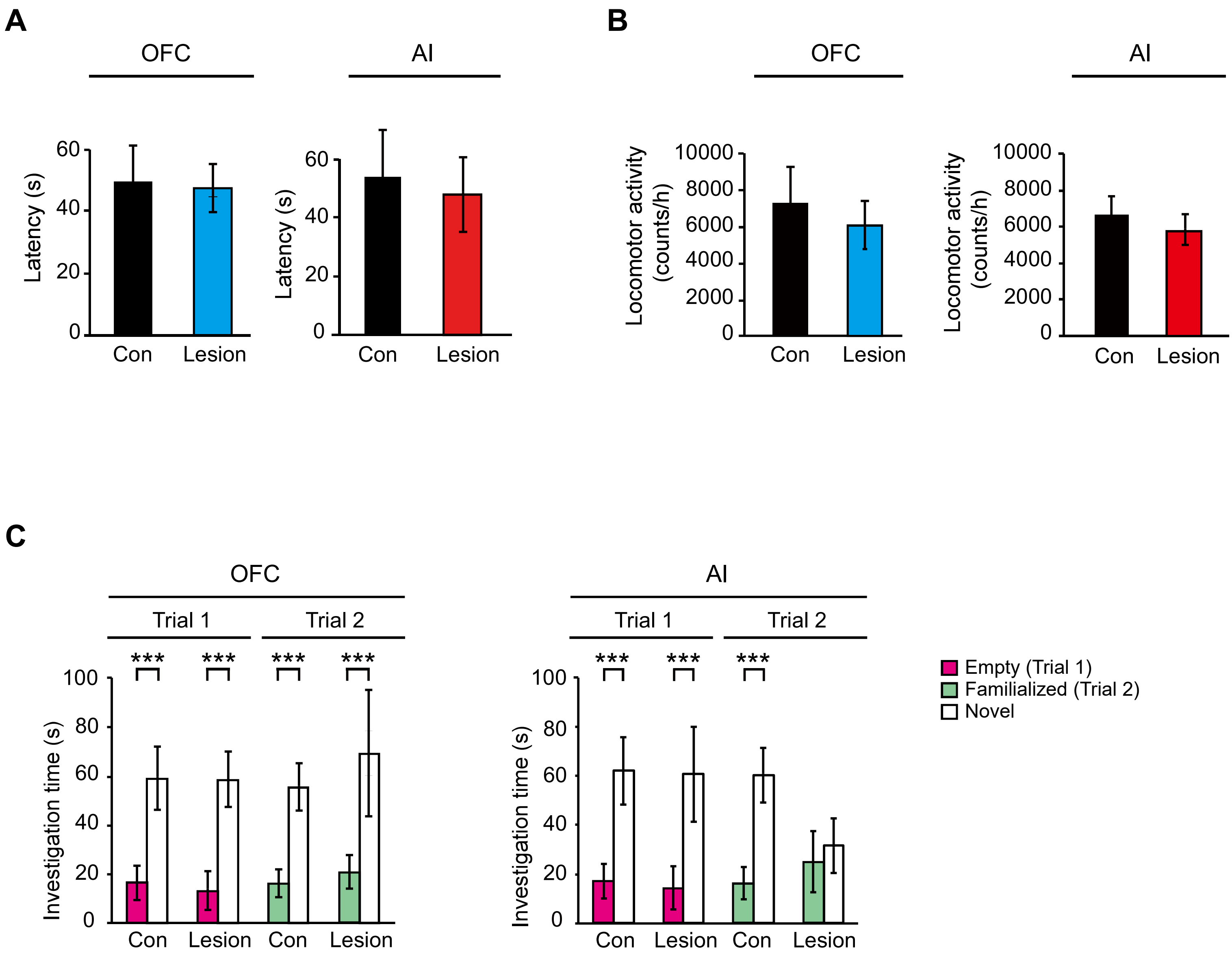
A
OFC
AI
Latency (s)
0
20
40
60
Con
Lesion
B
OFC
AI
Locomotor activity (counts/h)
0
2000
4000
6000
8000
10000
Con
Lesion
C
OFC
AI
Trial 1
Trial 2
***
Investigation time (s)
0
20
40
60
80
100
Con
Lesion
Empty (Trial 1)
Familialized (Trial 2)
Novel

### FIGURE S3

**A**

| Symbol | GWAS analysis | Class |
|---|---|---|
| Atat1 | 1.24 (22.0) | 5 |
| Flot1 | 1.24 (22.0) | 7 |
| Hist1h2bc | 1.21 (23.1) | 3 |
| Hist1h2bg | 1.21 (23.1) | 3 |
| Clic1 | 1.16 (19.1) | 1 |
| Hspa1l | 1.16 (19.1) | 7 |
| Ltb | 1.16 (19.1) | 1 |
| Usmg5 | 1.14 (13.4) | 2 |
| Actr1a | 1.12 (9.05) | 5 |
| Cyp26b1 | 1.11 (8.15) | 6 |
| Fyn | 1.11 (7.3) | 4 |
| Grm3 | 1.1 (9.52) | 1 |
| Clu | 1.1 (8.52) | 7 |
| Ptk2b | 1.1 (8.52) | 4 |
| Mef2c | 1.1 (8.3) | 3 |
| Ckb | 1.1 (8.22) | 6 |
| Igsf9b | 1.1 (12.05) | 2 |
| Sdccag8 | 1.09 (7.52) | 5 |
| Arl3 | 1.09 (19.4) | 4 |
| Cnnm2 | 1.09 (19.4) | 1 |
| Rangap1 | 1.09 (12.0) | 4 |
| Tcf19 | 1.09 (11.0) | 3 |
| Ank3 | 1.08 (8.52) | 5 |
| Slc7a6 | 1.08 (7.7) | 1 |
| Tcf4 | 1.08 (11.3) | 3 |
| Sbno1 | 1.08 (11.1) | 3 |
| Atp2a2 | 1.07 (8.52) | 1 |
| Ptn | 1.07 (8.52) | 7 |
| Tob2 | 1.07 (8.05) | 3 |
| Msi2 | 1.07 (7.7) | 3 |
| Mapk3 | 1.07 (12.1) | 4 |
| Sez6l2 | 1.07 (12.1) | 1 |
| Med8 | 1.07 (12.0) | 3 |
| St3gal3 | 1.07 (12.0) | 7 |
| Atp6v0b | 1.06 (9.4) | 1 |
| Phf1 | 1.06 (9.1) | 3 |
| Egr1 | 1.06 (9.0) | 3 |
| Ywhae | 1.06 (9.0) | 4 |
| Plcl1 | 1.06 (8.15) | 4 |
| Foxp1 | 1.06 (8.0) | 3 |
| Tle3 | 1.06 (7.7) | 3 |
| Ptp4a1 | 1.06 (7.52) | 4 |
| Ctnnd1 | 1.06 (7.3) | 2 |
| Tmx2 | 1.06 (7.3) | 7 |
| Bap1 | 1.05 (7.4) | 7 |
| Twf2 | 1.05 (7.4) | 5 |
| Cdh11 | 1.05 (7.3) | 2 |

**B**

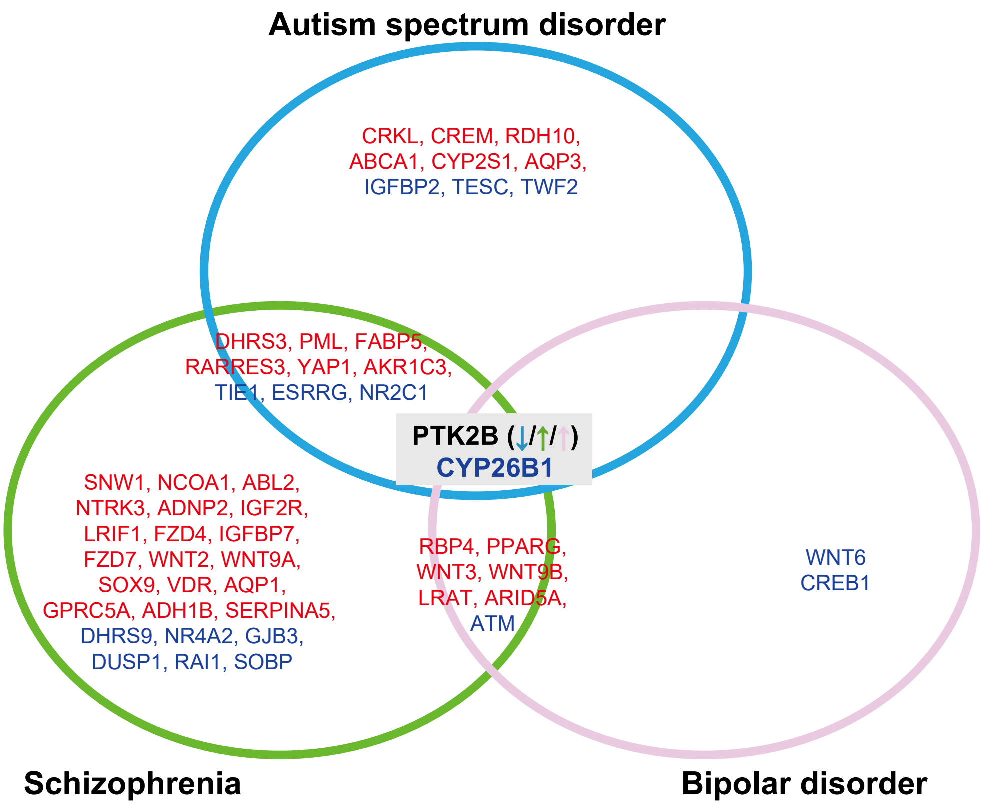

### FIGURE S4

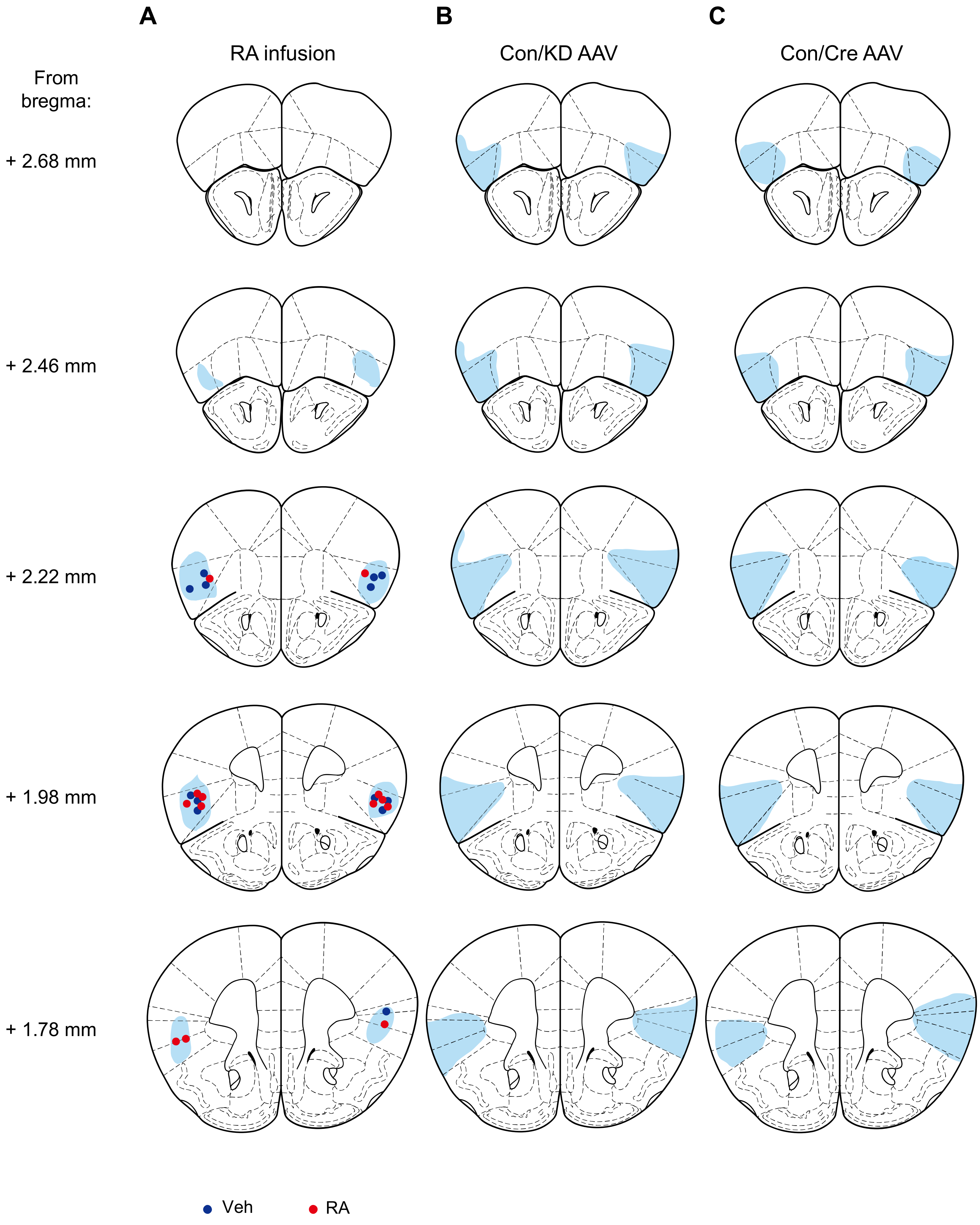
A
RA infusion
B
Con/KD AAV
C
Con/Cre AAV
From bregma:
+ 2.68 mm
+ 2.46 mm
+ 2.22 mm
+ 1.98 mm
+ 1.78 mm
Veh
RA

### FIGURE S5

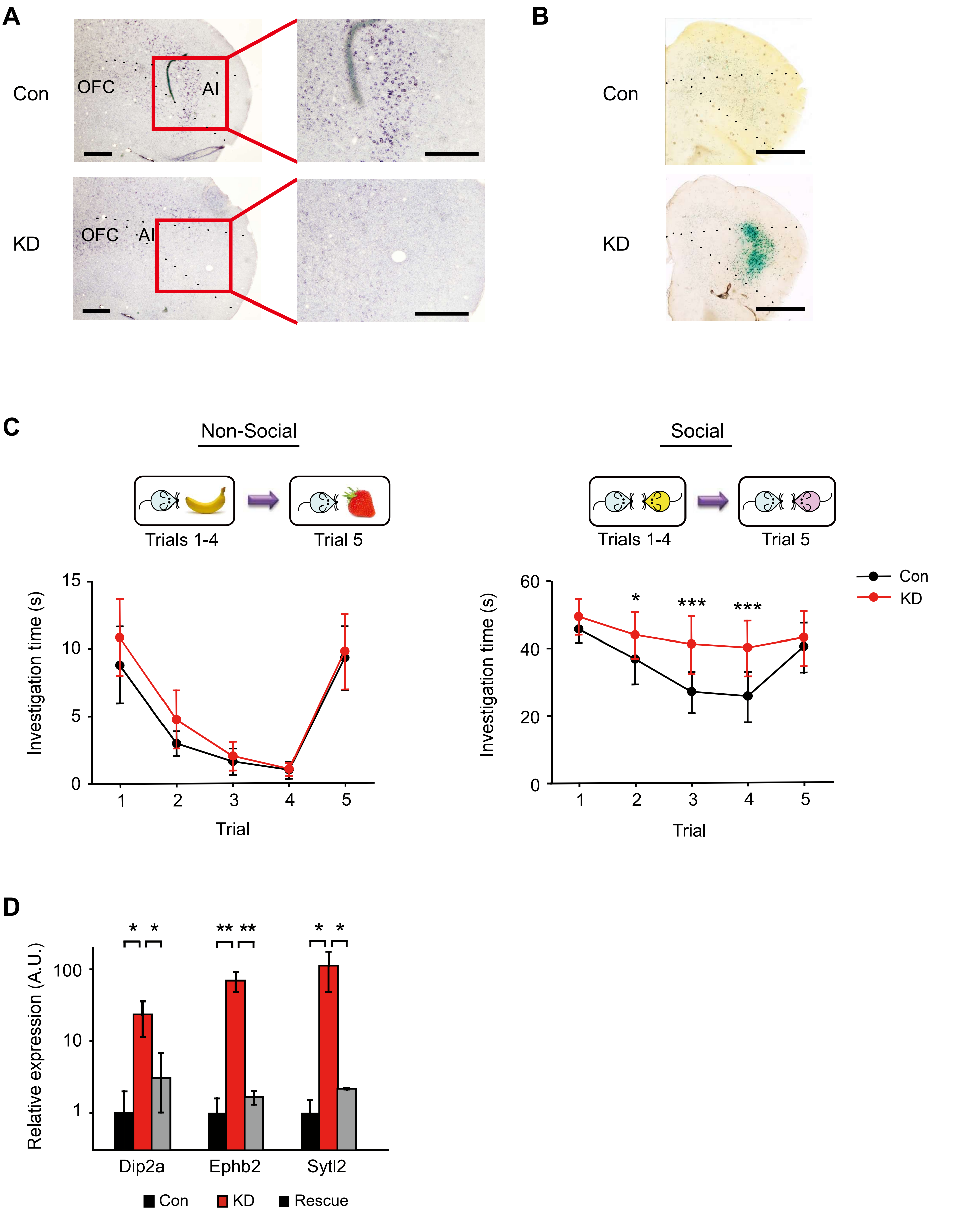

A
Con
OFC
AI
KD
OFC
AI
B
Con
KD
C
Non-Social
Trials 1-4
Trial 5
Social
Trials 1-4
Trial 5
Investigation time (s)
15
10
5
0
1
2
3
4
5
Trial
Investigation time (s)
60
40
20
0
*
***
***
1
2
3
4
5
Trial
Con
KD
D
*
*
**
**
*
*
Relative expression (A.U.)
100
10
1
Dip2a
Ephb2
Sytl2
Con
KD
Rescue

### FIGURE S6

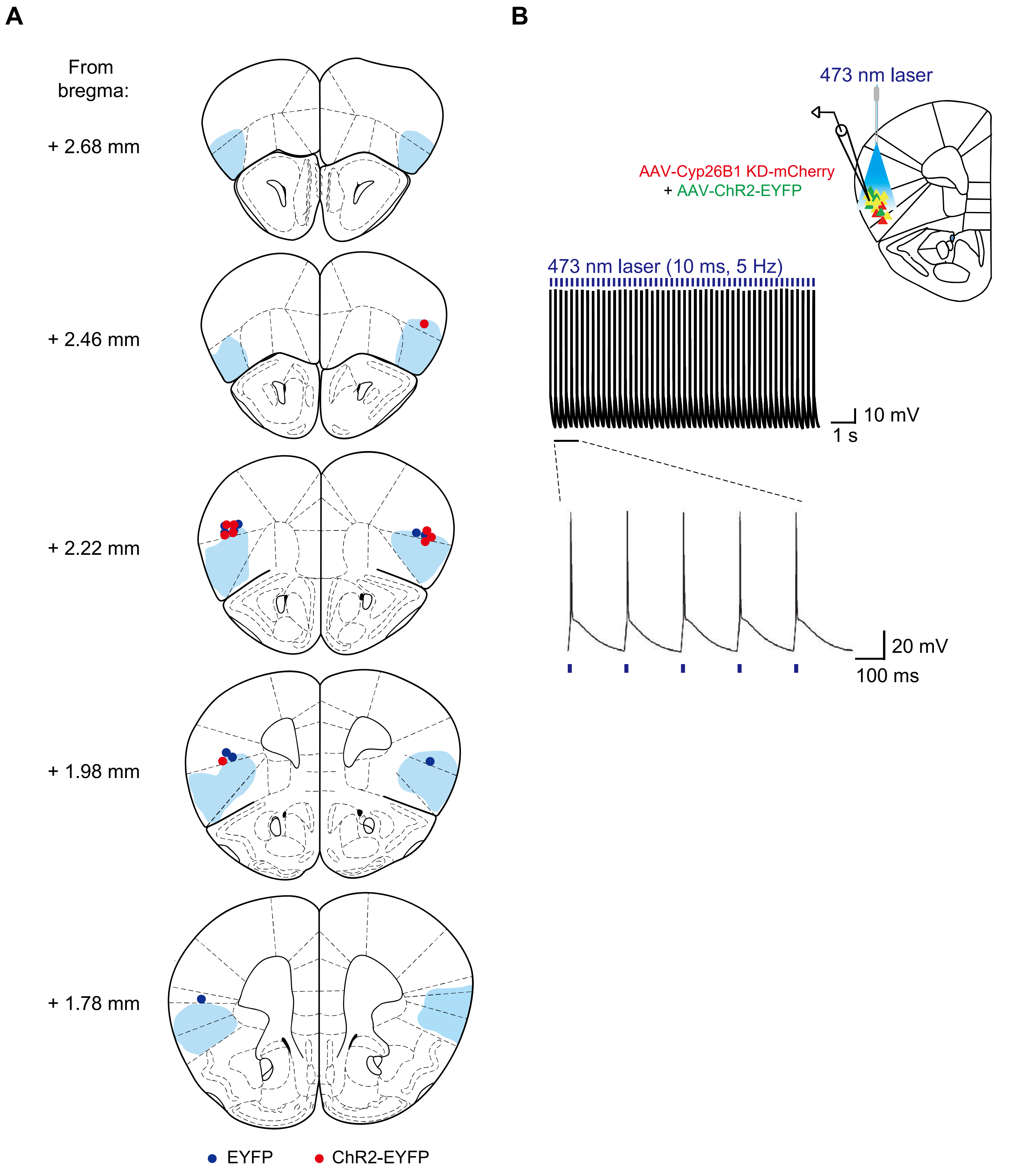
A
From bregma:
+ 2.68 mm
+ 2.46 mm
+ 2.22 mm
+ 1.98 mm
+ 1.78 mm
EYFP
ChR2-EYFP
B
473 nm laser
AAV-Cyp26B1 KD-mCherry
+ AAV-ChR2-EYFP
473 nm laser (10 ms, 5 Hz)
10 mV
1 s
20 mV
100 ms

FIGURE S7

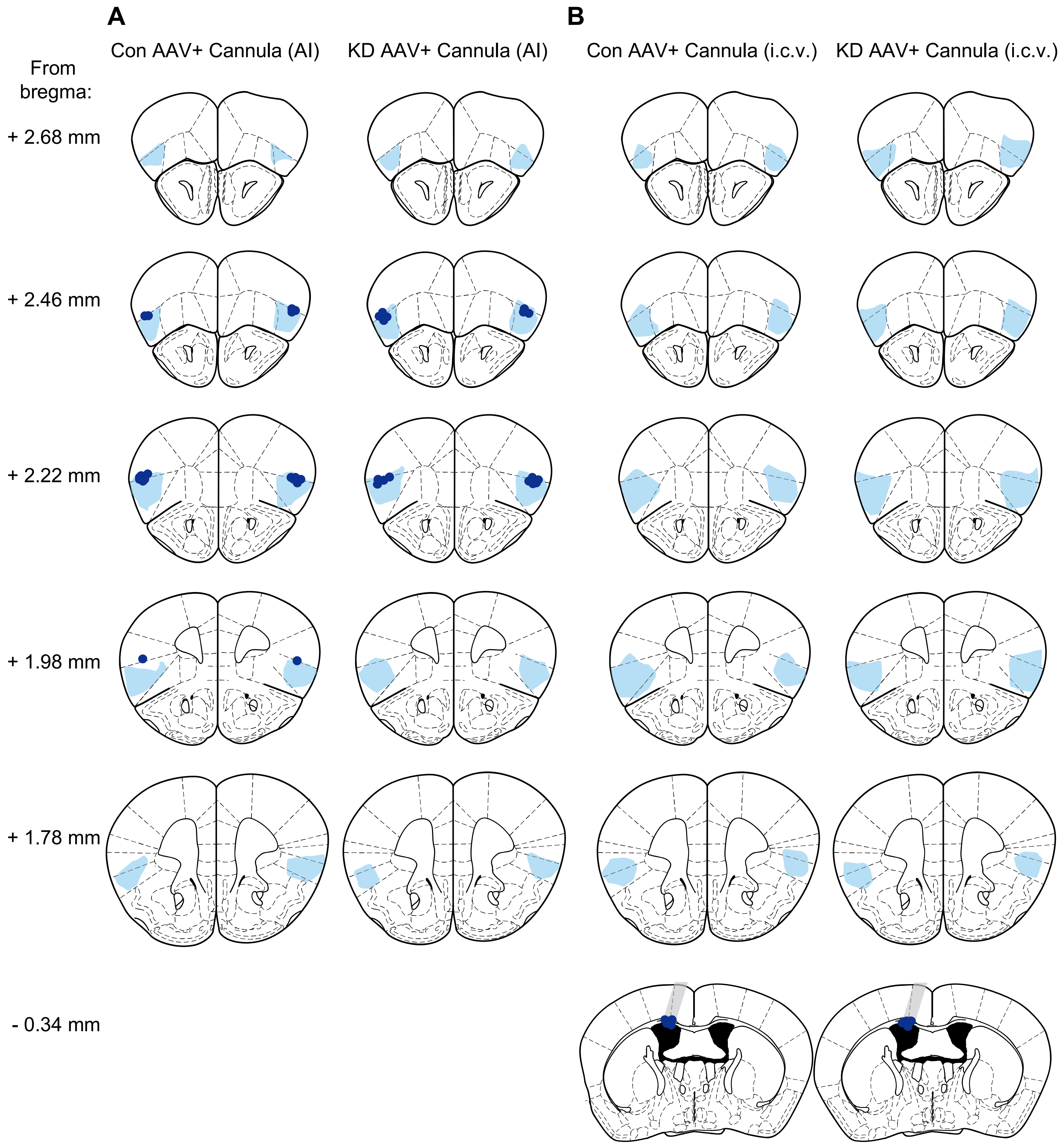
A
B
Con AAV+ Cannula (AI)
KD AAV+ Cannula (AI)
Con AAV+ Cannula (i.c.v.)
KD AAV+ Cannula (i.c.v.)
From bregma:
+ 2.68 mm
+ 2.46 mm
+ 2.22 mm
+ 1.98 mm
+ 1.78 mm
- 0.34 mm

**A**

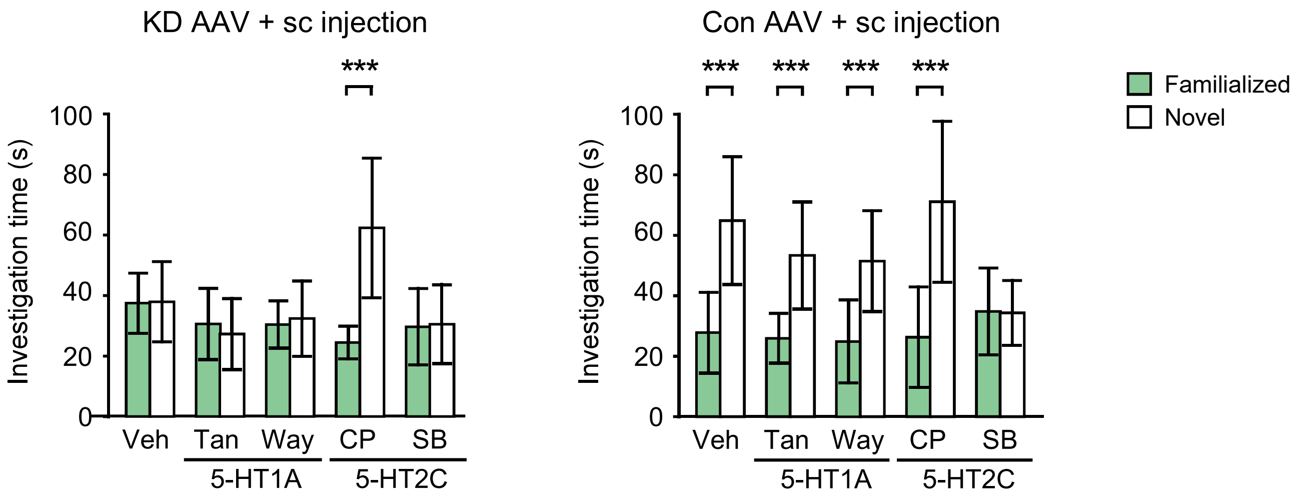
KD AAV + sc injection
Con AAV + sc injection
***
Investigation time (s)
0
20
40
60
80
100
Veh
Tan
Way
CP
SB
5-HT1A
5-HT2C
Familialized
Novel

**B**

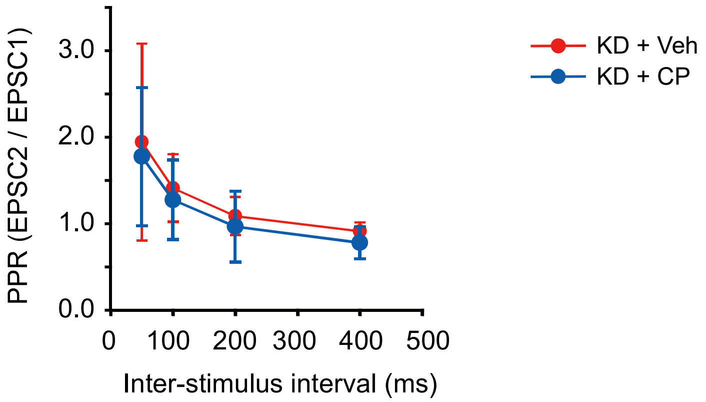
PPR (EPSC2 / EPSC1)
0.0
1.0
2.0
3.0
0
100
200
300
400
500
Inter-stimulus interval (ms)
KD + Veh
KD + CP

### FIGURE S9

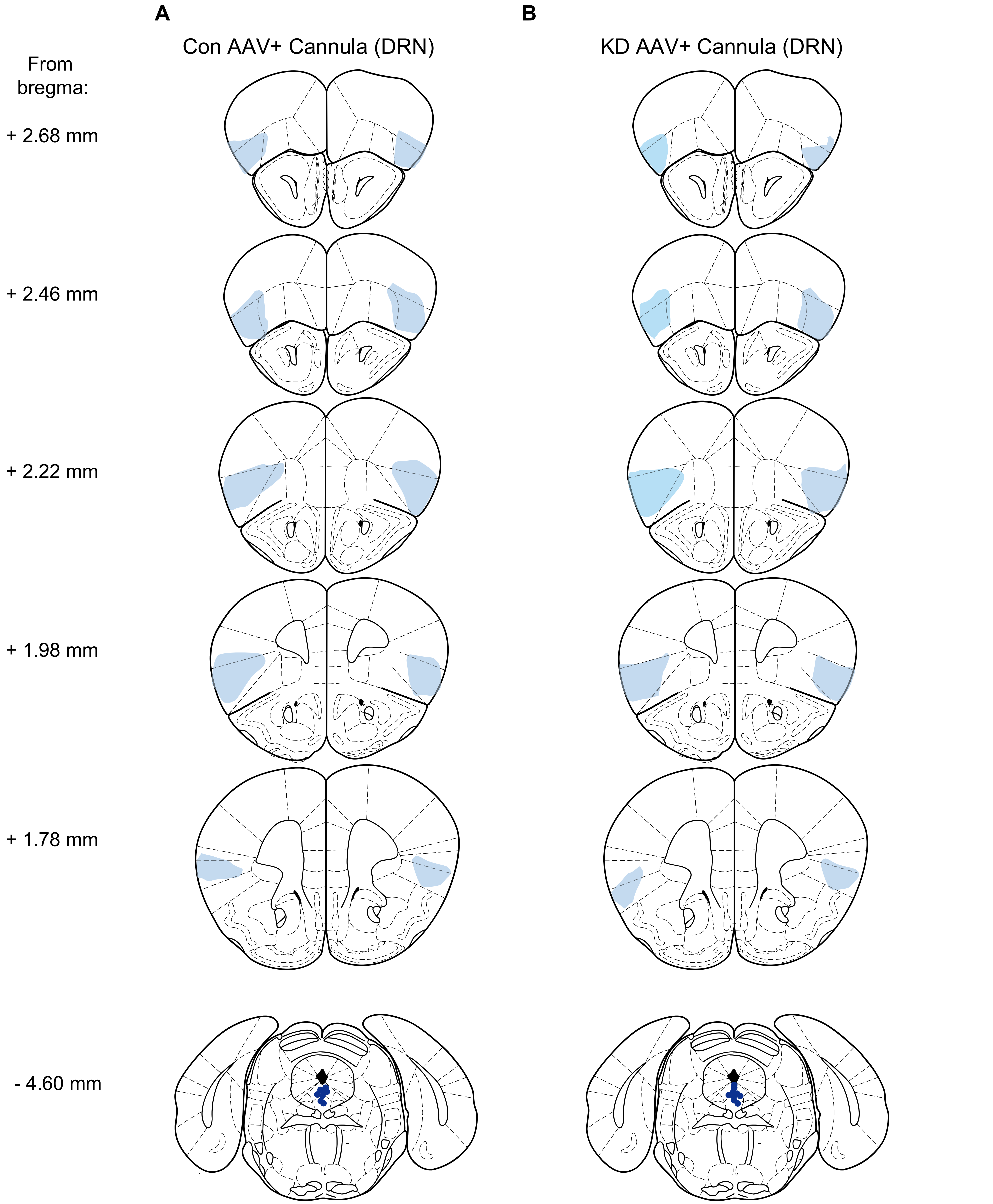

A
B
Con AAV+ Cannula (DRN)
KD AAV+ Cannula (DRN)
From bregma:
+ 2.68 mm
+ 2.46 mm
+ 2.22 mm
+ 1.98 mm
+ 1.78 mm
- 4.60 mm

**TABLE S1. List of genes differentially expressed between groups after Cyp26B1 control, KD, and rescue AAV infection**

| Gene | | Fold change | | Relevance to RA signaling | |
|---|---|---|---|---|---|
| **Symbol** | **Full Name** | **KD vs. Con** | **KD vs. Rescue** | **RARE** (*25*) | **RAR Binding** (*26*) |
| Scimp | SLP adaptor and CSK interacting membrane protein | -5.2 | -4 | | |
| Sp100 | nuclear antigen Sp100 | -2.2 | -2.1 | Yes | |
| Atad2 | ATPase family, AAA domain containing 2 | -1.7 | -1.6 | Yes | Yes |
| Samhd1 | SAM domain and HD | -1.7 | -1.9 | Yes | Yes |
| Treml2 | triggering receptor expressed on myeloid cells-like 2 | -2.6 | -3.6 | | |
| Zbp1 | Z-DNA binding protein 1 | -3.3 | -4.9 | | Yes |
| Oas2 | 2'-5' oligoadenylate synthetase 2 | -3.2 | -4.1 | Yes | Yes |
| Lat2 | linker for activation of T cells family, member 2 | -1.5 | -1.3 | | Yes |
| Pilra | paired immunoglobin-like type 2 receptor alpha | -2.3 | -1.6 | | |
| Map7 | microtubule-associated protein 7 | 2.8 | 7.4 | | |
| Serinc5 | serine incorporator 5 | 4.7 | 10.7 | | Yes |
| Sytl2 | synaptotagmin-like 2 | 5.4 | 11.8 | Yes | |
| Rabgap1l | RAB GTPase activating protein 1-like | 1.4 | 1.6 | Yes | Yes |
| Dip2a | disco interacting protein 2 homolog A | 3.4 | 5.5 | | Yes |
| Ctsk | cathepsin K | 1.5 | 1.6 | Yes | Yes |
| Ephb2 | Eph receptor B2 | 2.6 | 2.5 | Yes | Yes |
| Clcn2 | chloride channel, voltage-sensitive 2 | 2.4 | 2.5 | | Yes |
| Qdpr | quinoid dihydropteridine reductase | 2.4 | 2.6 | Yes | |
| Frmd5 | FERM domain containing 5 | 3.7 | 3.8 | Yes | Yes |
| Trim32 | tripartite motif-containing 32 | 1.9 | 1.6 | | |

Abbreviations: RA, retinoic acid; KD, knockdown; RARE, retinoic acid response element; RAR, retinoic acid receptor
